# Structure-Based Immunoinformatics Design of a CTB-Adjuvanted Multi-Epitope Mucosal Vaccine Against Helicobacter pylori

**DOI:** 10.64898/2026.06.16.732557

**Authors:** Razieh Veisi, Amin Mohsenzadeh, Nahal Hadi, Raham Armand

**Author notes:** **Corresponding authors**: Razieh Veisi; Nahal Hadi.

## Abstract

**Background:** Helicobacter pylori coloniz the gastric mucosa of nearly half of the global population and is classified as a Group I carcinogen by the World Health Organization due to its strong association with gastric cancer. The growing prevalence of antibiotic-resistant H. pylori strains significantly compromises current therapeutic strategies, emphasizing the urgent need for effective prophylactic approaches.

**Research design and methods:** In this study, a novel multi-epitope vaccine was designed targeting H. pylori, incorporating epitopes from four key virulence proteins: BabB, SabB, SabA, and VacA. Using an immunoinformatics-guided structural vaccinology approach, B- and T-cell epitopes were predicted, prioritized based on immunogenicity, conservation, population coverage, and non-homology to human proteins, and assembled into the final vaccine construct. To enhance immunogenicity and specifically stimulate mucosal immune responses, the cholera toxin B subunit (CTB) was fused at the N-terminal via an EAAAK linker, a novel application in H. pylori multi-epitope vaccines. The PADRE universal epitope and additional linkers were incorporated to optimize epitope presentation and helper T-cell activation.

**Results:** Comprehensive evaluations of physicochemical, antigenic, allergenic, and toxic properties were conducted, followed by secondary and tertiary structure modeling, refinement, and validation. Conformational B-cell epitopes were mapped, and molecular docking, binding affinity analysis, energy minimization, and molecular dynamics simulations confirmed structural stability and re-ceptor interactions. Codon optimization and in silico cloning predicted efficient expression in Escherichia coli, while immune simulations suggested robust humoral and cellular responses.

**Conclusions:** This study presents a promising multi-epitope vaccine candidate against H. pylori, offering a rational framework for future experimental validation and potential clinical application.

## Background

*Helicobacter pylori* is a highly adaptive, Gram-negative bacterium that colonizes the human gastric mucosa and is responsible for one of the most persistent bacterial infections worldwide (1). It is estimated that more than half of the global population carries *H. pylori*, with a particularly high prevalence in developing countries. Its chronic colonization is strongly associated with peptic ulcer disease, atrophic gastritis, mucosa-associated lymphoid tissue (MALT) lymphoma, and gastric adenocarcinoma, which is classified by the World Health Organization as a Group I carcinogen (2,3). Despite decades of research, the ability of *H. pylori* to evade immune responses, establish lifelong infection, and develop resistance to antibiotics has made its control increasingly challenging.

The growing emergence of antibiotic-resistant *H. pylori* strains has severely limited the effectiveness of standard triple and quadruple therapies. Resistance to clarithromycin, levofloxacin, and metronidazole has increased dramatically across many regions, contributing to treatment failure rates exceeding 20-40% in certain populations (4,5). These challenges underscore the urgent need for alternative preventive strategies, particularly the development of safe, effective, and widely applicable vaccines capable of inducing long-lasting immunity at the gastric mucosal surface, the primary site of infection (6). Reverse vaccinology and immunoinformatics have emerged as transformative approaches that allow the systematic identification of protective antigens and immunogenic epitopes directly from pathogen genomes (7). These computational strategies significantly reduce the time, cost, and experimental complexity associated with conventional vaccine development, while enabling the precise selection of highly conserved, antigenic, and non-allergenic epitopes (8–10). In the context of *H. pylori*, outer membrane-associated virulence proteins such as BabB, SabA, SabB, and VacA represent promising vaccine targets due to their established roles in adhesion, colonization, and immune modulation. Their high surface exposure and conservation across strains make them attractive candidates for designing broad-spectrum vaccines.

In multi-epitope vaccine design, combining carefully screened B-cell, Cytotoxic T-lymphocyte (CTL), and Helper T-lymphocyte (HTL) epitopes allows the generation of vaccines that are capable of eliciting both humoral and cellular immune responses (11–13). This is particularly relevant for *H. pylori*, where both arms of the immune system are required to restrict colonization and prevent chronic persistence (14). In this study, the *H. pylori* pan-proteome was systematically analyzed, followed by epitope prediction, selection of fully conserved regions, population coverage assessment, allergenicity and toxicity evaluation, structural modeling, molecular docking, molecular dynamics simulation, immune simulation, and in-silico cloning. Collectively, these steps enabled the rational design of a stable and immunogenic multi-epitope vaccine construct. A major innovation of the present research is the incorporation of the cholera toxin B subunit (CTB) as a mucosal adjuvant at the N-terminal of the vaccine construct. Although CTB has been examined as an effective immunomodulatory component in several viral and bacterial vaccines, its incorporation into multi-epitope vaccine constructs targeting *H. pylori* has not been previously reported (15–17). CTB naturally binds with high affinity to GM1 ganglioside receptors on the surface of intestinal epithelial cells, facilitating efficient uptake and presentation of fused antigens (18). While the molecular details of its interaction with innate immune receptors were not the focus of the present study, CTB is well-recognized for its capacity to enhance antigen-specific responses and promote strong mucosal immunity, an essential feature for combating pathogens that establish infection within the gastrointestinal tract (19). Computational validation through docking, MD simulation, and immune simulation demonstrated strong interaction with immune receptors and the ability to induce robust humoral and cellular responses. Taken together, this study presents a comprehensive computational framework for the development of a novel CTB-adjuvanted multi-epitope vaccine targeting *H. pylori* Figure 1.

**Figure 1.**
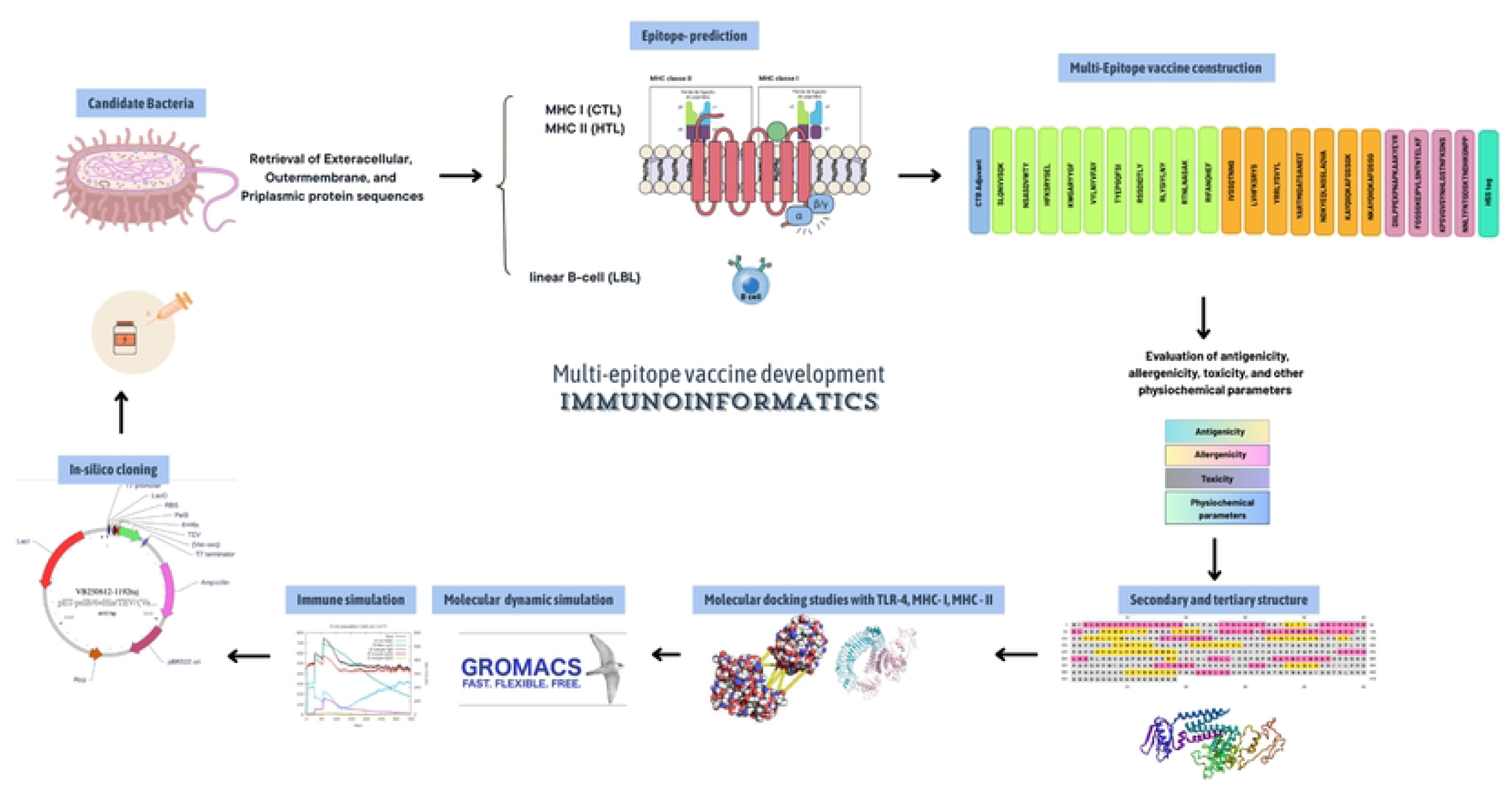
Schematic workflow of in silico multi-epitope vaccine design process. (Schematic illustration generated using Canva (Version 1.121.0)) By integrating reverse vaccinology, immunoinformatics, structural modeling, and immune simulation, the findings provide a strong foundation for future in-vitro and in-vivo validation. Ultimately, this approach offers a promising pathway toward an effective vaccine capable of reducing the global burden of *H. pylori* infection.

## Methods

This study employed multiple web servers and bioinformatics tools for in silico analyses; a complete list is provided in Supplementary Table S1.

### Retrieval of *Helicobacter pylori* pan-proteome and antigenicity prediction of proteins

In the first step, the complete sequences of selected structural and virulence-associated proteins from the *Helicobacter pylori* pan-proteome were retrieved from the UniProtKB/Swiss-Prot database (https://www.uniprot.org/). Four proteins exhibiting high antigenic potential were identified and selected for further analysis Supplementary Table S2. The accuracy and annotation of these sequences were verified against UniProtKB/Swiss-Prot entries to ensure reliability. The antigenicity of the selected proteins was predicted using ANTIGENpro (https://scratch.proteomics.ics.uci.edu/) and VaxiJen v2.0 (http://www.ddg-pharmfac.net/vaxijen/VaxiJen/VaxiJen.html). Proteins with ANTIGENpro scores above 0.7 (20) and VaxiJen scores above 0.4 (21) for the bacterial category were considered highly antigenic and were retained for subsequent epitope prediction. Additionally, the conservation, human homology, allergenicity, and toxicity of the selected proteins were evaluated. Following this screening, predicted cytotoxic T lymphocyte (CTL), helper T lymphocyte (HTL), and linear B lymphocyte (LBL) epitopes derived from these proteins were compiled into an epitope repository for multi-epitope vaccine design.

### Prediction of T-cell epitopes

Cytotoxic T lymphocyte (CTL) epitopes were identified using the NetCTL 1.2 server (22) (https://www.cbs.dtu.dk/services/NetCTL/) in combination with the Immune Epitope Database (IEDB) MHC I prediction tools (23) (https://tools.iedb.org/mhci/). The NetCTL server predicts CTL epitopes with high accuracy, exhibiting a specificity of 94–99% and a sensitivity between 54% and 89% (24). In this study, the four selected protein sequences were screened against the most common HLA class I alleles (A1, A2, A3, A24, A26, B7, B8, B27, B39, B44, B58, and B62) to predict and assess potential cytotoxic T lymphocyte (CTL) epitopes (25). Additionally, an extended set of HLA class I-restricted epitopes was predicted using the stabilized matrix method (SMM) implemented in the IEDB MHC I binding tools. Peptides with a percentile rank ≤ 2, an IC₅₀ value below 200 nM, and high binding scores were classified as strong binders (26). CTL epitopes predicted across multiple alleles by both the NetCTL 1.2 and IEDB servers were compared, and overlapping high-affinity epitopes meeting the desired thresholds were selected for downstream analyses. For helper T lymphocyte (HTL) epitope prediction, 15-mer peptides were screened against a comprehensive set of HLA class II alleles, including HLA-DR, HLA-DP, and HLA-DQ, using the NetMHCIIpan 3.2 server (27). Based on percentile rank thresholds of ≤2%, 2–10%, and >10%, the predicted peptides were categorized as strong, moderate, and non-binders, respectively (28). Using the SMM-align method (NetMHCII 1.1) available in the IEDB server (29), an additional set of HTL epitopes of the same length was predicted across 54 HLA class II alleles, encompassing HLA-DR, HLA-DQ, and HLA-DP allele groups (23). Epitopes exhibiting IC50 < 200 nM, a percentile rank ≤ 2, and predicted as strong binders by multiple alleles across both methods were selected for further evaluation. The IEDB platform predicts peptide binding to each MHC-II molecule using artificial neural networks (ANN) trained on a dataset comprising over 500,000 binding affinity (BA) and eluted ligand mass spectrometry (EL) measurements, ensuring high predictive reliability (26). For subsequent analyses, all selected CTL and HTL epitopes were evaluated through a series of filtering criteria, including immunogenicity, antigenicity, allergenicity, and toxicity. These properties were predicted using the IEDB Class I Immunogenicity server, VaxiJen v2.0, AllerTOP 2.0, and ToxinPred, respectively.

### Prediction of LBL epitopes

In this study, linear (continuous) B-cell epitopes were predicted using default parameters across five servers: ABCPred (30), BCPreds (31), BepiPred (32), and SVMtrip (33). To enhance prediction accuracy, ABCPred (http://www.imtech.res.in/raghava/abcpred/) employs a recurrent neural network to differentiate epitopes from non-epitopes (30). BCPreds (http://ailab.ist.psu.edu/bcpred/) utilizes kernel-based methods and a support vector machine (SVM) model, achieving an area under the curve (AUC) value of 0.758 (31). Additionally, IEDB Emini (34) provides access to surface accessibility data at the B-cell level, aiding in epitope identification. The BepiPred-2.0 web server (http://www.cbs.dtu.dk/services/BepiPred/) employs a random forest algorithm trained on a large dataset of linear B-cell epitopes obtained from the IEDB database (32). The SVMTrip server (http://sysbio.unl.edu/SVMTriP/) applies a support vector machine to combine sequence similarity, tri-peptide affinity, and antigenic epitope prediction, achieving an AUC of 0.702 (33). Initially, BepiPred was employed for primary prediction of linear B-cell epitopes, and the results were cross-validated using four additional servers. A predicted epitope was considered valid only if it was confirmed by at least one of the other servers. Assessing the surface accessibility of epitopes from structural proteins is essential to ensure their localization in solvent-exposed regions suitable for B-cell recognition (35), which was evaluated using the Emini tool from IEDB (34,36). Consequently, epitopes not located on the protein surface were excluded. The final set of linear B-cell epitopes on the surface of structural proteins was selected based on their antigenicity, non-allergenicity, and non-toxicity.

### Multiple alignment and selection of fully conserved epitopes

Following multiple alignments of the desired structural protein sequences for various strains using clustalOmega (37), the modifications made, such as deletions, additions, and substitutions, were identified in each strain (38). Then, the predicted CTL, HTL, and LBL epitopes were compared with the results of this alignment, and epitopes that remained constant in all strains were selected as fully conserved epitopes for inclusion in the final vaccine construct (38).

### Population coverage, epitope protection analysis, and autoimmunity identification

The IEDB population coverage analysis tool (https://tools.iedb.org/population/) was used with default settings to evaluate the immune responsiveness of the predicted CTL and HTL epitopes, considering their corresponding HLA genotype frequencies and ensuring sufficient coverage across the global human population (39). To evaluate the degree of conservation or variability of the selected T- and B-cell epitopes, BLASTp (https://blast.ncbi.nlm.nih.gov/Blast.cgi) was used to compare these sequences against related proteins. This analysis ensured that the chosen epitopes exhibit optimal conservation across similar sequences, supporting their suitability for vaccine design (40). The degree of conservation of each epitope was assessed using BLASTp (https://blast.ncbi.nlm.nih.gov/Blast.cgi), which provides conservation scores ranging from 0% (minimum) to 100% (maximum) (41). To prevent potential host autoimmune responses and cross-reactivity, all selected epitopes were further compared against the human proteome in the UniProt database using BLASTp. Any epitope exhibiting ≥35% sequence identity with human proteins was considered homologous and excluded from vaccine design. Consequently, only non-homologous epitopes were retained for the construction of the multi-epitope vaccine (42,43).

### Multi-epitope vaccine design

In this study, an overlapping strategy was applied to avoid the inclusion of redundant epitopes during the selection of suitable candidates for vaccine construction. Epitopes were prioritized based on their immunogenicity, non-allergenicity, high population coverage, absence of similarity to human proteins, and surface accessibility, and subsequently assembled into the final design of the multi-epitope vaccine (44). The final multi-epitope vaccine construct comprises 486 amino acid residues, including Adjuvant (cholera toxin B subunit/CTB (MIKLKFGVFFTVLLSSAYAHGTPQNITDLCAEYHNTQIYTLNDKIFSYTESLAGKREMAIITFKNGAIFQVEVPGSQHIDSQKKAIERMKDTLRIAYLTEAKVEKLCVWNNKTPHAIAAISMAN), the PADRE universal epitope, 10 CTL epitopes, 7 HTL epitopes, 4 linear B-cell epitopes. Within the vaccine design, adjuvant and PADRE were connected using EAAAK linker, CTL epitopes were linked using AAY (Ala-Ala-Tyr) linkers, HTL epitopes were joined via GPGPG (Gly-Pro-Gly-Pro-Gly) linkers, LBL epitopes were linked by KK (Lys-Lys) linkers, and LBL epitopes were linked to H5E tag using GGGGS linker (45–47).

PADRE, which has the amino acid sequence AKFVAAWTLKAAA, is a simple carrier epitope. Combined with an adjuvant, it provides an effective response and can be used in making multi-epitope and recombinant vaccines (48). To activate helper T-cells (CD4+ T-cells) and reduce the polymorphism of HLA-DR molecules in the population, this epitope has been added to the vaccine structure with the GPGPG linker. In addition, to further enhance and prolong the immune response, the cholera toxin B subunit (CTB) adjuvant was incorporated at the N-terminal of the vaccine construct through the EAAAK linker. This placement at the N-terminal region facilitates effective antigen presentation and improves the overall immunogenicity of the multi-epitope vaccine (49). Given that the Food and Drug Administration (FDA) and the European Medicines Agency (EMA) advise against the use of the 6His tag due to its potential to elicit undesirable immune responses, alternative purification tags have been explored. Humanized tags derived from human BLAST proteins, such as H3A (HAAHAH), H5T (HTHTHTHTH), and H5E (HEHEHEHEH), have been developed and evaluated, demonstrating promising performance in both purification efficiency and functional behavior within human biopharmaceutical applications (50). Consequently, in this construct, the H5E tag was incorporated at the C-terminal end to facilitate protein purification and detection.

### Evaluation of antigenicity, allergenicity, toxicity, solubility and physicochemical properties

To evaluate the antigenic potential of the designed vaccine, the VaxiJen v2.0 server (http://www.ddg-pharmfac.net/vaxijen/VaxiJen/VaxiJen.html) was employed using the bacterial model and a threshold of 0.4. VaxiJen predicts antigenicity by converting protein sequences into consistent numerical vectors through the auto-cross covariance (ACC) method, which captures the intrinsic physicochemical features of amino acids (51). Additionally, the AntigenPro server was utilized as a complementary predictor. AntigenPro is a sequence-based, alignment-free, and pathogen-independent tool that estimates protein antigenicity using machine learning-derived models (51). In this study, antigenicity was confirmed based on the following criteria: VaxiJen (bacterial model) scores above 0.4 and ANTIGENpro scores exceeding 0.7. Allergenicity was assessed using AllergenFP 1.0 (http://ddg-pharmfac.net/AllergenFP/) and AllerTOP 2.0 (https://www.ddg-pharmfac.net/AllerTOP/). AllergenFP applies a binary classification system to differentiate allergens from non-allergens, while both tools employ the automatic cross-covariance (ACC) method, converting amino acid strings into standardized feature vectors after encoding them with E-descriptors (52,53). Additionally, ToxinPred was used to predict the toxicity of the vaccine construct and its individual epitopes. This server employs the SVM model and a dataset of 1,805 toxic peptides (<35 residues) (54)to distinguish toxic sequences from non-toxic ones. Solubility was evaluated using both Protein-sol (https://protein-sol.manchester.ac.uk) and SolPro (http://scratch.proteomics.ics.uci.edu). Unlike SolPro, which relies on an SVM-based algorithm and achieves an overall accuracy of more than 74% under tenfold cross-validation (55), Protein-sol predicts solubility using empirical data derived from *E. coli* proteins expressed in a cell-free system (56). Several physicochemical parameters of the final vaccine construct, including molecular weight, theoretical isoelectric point (pI), charge, extinction coefficient, estimated half-life, instability index, aliphatic index, and GRAVY score (57), were calculated using the ExPASy ProtParam tool (https://web.expasy.org/protparam/). Furthermore, SignalP 4.1 (58)(https://www.cbs.dtu.dk/services/SignalP/) and TMHMM v2.0 (59) (https://www.cbs.dtu.dk/services/TMHMM/) were employed to detect the potential presence of signal peptides and transmembrane helices, respectively, within the vaccine sequence.

### Secondary structure prediction

The secondary structure of the vaccine construct includes several structural elements, such as alpha-helical regions, extended strands, beta sheets, and random coils. These features were evaluated using the SOPMA server (60) (https://npsa-prabi.ibcp.fr/cgi-bin/npsa_automat.pl?page=/NPSA/npsa_sopma.html) and PSIPRED v4.0 (61)(http://bioinf.cs.ucl.ac.uk/psipred/). Both tools provide secondary-structure predictions with an accuracy exceeding 80%. PSIPRED performs its analysis through two feed-forward neural networks that interpret the output of PSI-BLAST (Position-Specific Iterated BLAST) searches (61). After submitting the vaccine sequence to the SOPMA server, the output width parameter was adjusted to 70. The settings for secondary structure prediction were configured to include four conformational states (helix, sheet, turn, and coil), with a similarity threshold of 8 and a window width of 17, ensuring reliable structural profiling.

#### Modeling, refinement, and validation of 3D structure

Robetta (https://robetta.bakerlab.org/), a comprehensive online platform for protein structure prediction was utilized to model the final 3D conformation of the vaccine construct (62,63). This server performs continuous evaluation of multiple prediction criteria, including structural coverage, local accuracy, and model completeness. Robetta employs state-of-the-art deep-learning frameworks such as RoseTTAFold and TrRosetta, together with homology modeling and fragment-based assembly. Its interactive interface also supports user-defined sequence alignments, structural constraints, and fragment libraries, enabling rapid and highly accurate protein structure prediction (64). In the subsequent stage, the predicted 3D vaccine model was subjected to structural refinement using GalaxyRefine (65) (http://galaxy.seoklab.org/cgi-bin/submit.cgi?type=REFINE). This platform applies molecular dynamics-based procedures, beginning with rebuilding and repacking of side chains, followed by global relaxation steps that alleviate steric conflicts and enhance structural integrity. According to CASP10 evaluations, GalaxyRefine ranks among the most effective methods for improving local structural quality (66). The refined structure was then assessed using several standard metrics (67), including GDT-HA, RMSD, MolProbity score, clash score, and Ramachandran plot. For model validation, the structure was analyzed using the ProSA-web server (68) (https://prosa.services.came.sbg.ac.at/prosa.php) and SAVES v6.0 (https://saves.mbi.ucla.edu). ProSA determines both overall and residue-level quality based on the Z-score, providing an estimate of structural correctness. Deviations of the Z-score from the typical range of experimentally determined proteins indicate a higher likelihood of structural inaccuracies in the predicted model. A local quality score graph is displayed within the 3D molecular viewer (68) to facilitate the identification of regions with potential structural issues. To further assess the structural integrity of the predicted model, several validation tools integrated into the SAVES v6.0 platform, namely ERRAT (69), VERIFY3D (70), PROVE (66), PROCHECK (66), and WHATCHECK (71), were employed. These tools collectively examine different stereochemical and structural parameters of the protein. Among them, PROCHECK specifically analyzes stereochemical quality by evaluating the geometry of residues and overall conformational parameters of the model (72,73). Multiple servers are available for generating the Ramachandran plot, including PDBsum (74), MolProbity (75), STAN Server (76), and RAMPAGE (77). In the present study, the PDBsum server was used for this purpose. Finally, the refined vaccine model was visualized and inspected using Pymol, the final model of the vaccine structure was drawn(78).

### Prediction of conformational B-cell epitopes

After the 3D structure of the vaccine was modeled, refined, and validated, conformational (discontinuous) B-cell epitopes were identified using the ElliPro server (79) (http://tools.iedb.org/ellipro). ElliPro predicts conformational epitopes by assessing the protein’s geometric shape, solvent accessibility, and structural flexibility, typically generating longer epitope segments than linear predictors. In this analysis, the prediction thresholds were set to a minimum score of 0.8 and a maximum distance of 6 Å. ElliPro is considered a robust tool for mapping antibody-recognition regions, with an AUC performance of 0.732 (79).

### Molecular docking and binding affinity analysis

The 3D structures of TLR4, MHC-I, and MHC-II receptors were obtained from the Protein Data Bank (PDB) (https://www.rcsb.org) with the identifiers 3fxi, 1i4f, and 1aqd, respectively. All ligands and water molecules were removed from these structures prior to docking. The CPORT tool (Consensus Prediction of Interface Residues in Transient complexes) was employed to predict the active and passive residues involved in protein–protein interactions. Subsequently, the docking analysis was performed using the ClusPro server (https://cluspro.bu.edu/), which predicts protein–protein complexes based on rigid-body docking, scoring functions, and clustering of low-energy conformations. Based on the largest cluster size and lowest energy score provided by ClusPro, the best docking models for each complex were selected. The PRODIGY web server was then used to estimate the binding affinity of each docked complex by analyzing the top-ranked model from the best cluster. PRODIGY predicts binding affinities and thermodynamic parameters of protein–protein interactions using structural data from crystallography (27,80). The binding affinities of the four docked complexes were calculated at 37 °C and reported as Kd values, and the corresponding ΔG values (kcal mol⁻¹) were also obtained. Finally, the residues involved in interactions between the vaccine and the target receptors were visualized using PyMol and mapped with PDBsum (81).

### Energy minimization and molecular dynamics simulation

GROMACS v5.1, an open-source molecular dynamics (MD) simulation package running on Linux, was employed to perform energy minimization, equilibration, and production MD simulations of the designed multi-epitope vaccine ((82); Abraham et al., 2015). System topology files were generated using the OPLS-AA (Optimized Potential for Liquid Simulations–All Atom) force field (84). The vaccine construct was solvated in an octahedral box of SPC216 water molecules, with a minimum distance of 10 Å between any protein atom and the box boundary. Counterions were added to neutralize the system. Covalent bonds involving hydrogen atoms were constrained using the LINCS algorithm (85),, and long-range electrostatic interactions were calculated with the Particle Mesh Ewald (PME) method using a 10 Å real-space cutoff (38). Energy minimization was performed for 50,000 steps using the steepest-descent algorithm to remove steric clashes and unfavorable contacts. Subsequently, a two-phase equilibration protocol was applied: NVT ensemble: the system was gradually heated to 300 K over 100 ps under constant volume, with backbone restraints. NPT ensemble: 1 ns equilibration at 1 atm and 300 K without restraints, using the V-Rescale thermostat and Parrinello–Rahman barostat, with a 2 fs integration time step (Bussi et al., 2007).

Following equilibration, a 10 ns production MD run was carried out, with coordinates recorded every 10 ps. Trajectories were analyzed using standard GROMACS tools, and graphical outputs were generated with Xmgrace (87). Backbone stability of the vaccine construct was evaluated via Root Mean Square Deviation (RMSD). The RMSD increased initially from ∼0.02 nm to ∼0.9 nm and reached a plateau, indicating that the system attained structural equilibrium. Residue-level flexibility was assessed through Root Mean Square Fluctuation (RMSF), showing that most residues remained stable (0.02–0.04 nm), while surface-exposed loops exhibited slightly higher fluctuations (∼0.08 nm), which is consistent with natural protein flexibility. The radius of gyration (Rg) was calculated to monitor structural compactness, remaining nearly constant at 2.7–2.9 nm throughout the simulation, confirming that the vaccine construct maintained a stable and compact conformation. Overall, the MD simulation results demonstrate that the vaccine construct is structurally stable and suitable for subsequent molecular docking and immunoinformatics analyses.

### Codon optimization and in-silico simulation of vaccine

The Java Codon Adaptation Tool (JCat) (http://www.prodoric.de/JCat) was employed for codon optimization and reverse translation of the vaccine construct. JCat enables codon adaptation across a wide range of sequenced prokaryotic and eukaryotic hosts (87). Following optimization for *E. coli* strain K12, the corresponding cDNA sequence was generated to ensure efficient expression in this system. During the optimization process, additional parameters were set to prevent the introduction of rho-independent transcription terminators, prokaryotic ribosome-binding sites, and unwanted restriction enzyme recognition sites. The optimization results provided two key indicators of expression efficiency: the guanine-cytosine (GC) content and the codon adaptation index (CAI), both of which reflect the protein’s potential expression level in the selected host system (87). For subsequent in-silico cloning, recognition sites for the BamI and XhoI restriction enzymes were incorporated at the 5′ (N-terminal) and 3′ (C-terminal) ends of the optimized sequence, respectively. The finalized construct was then inserted into the pET-28a(+) expression vector using SnapGene 6.2 software (88) (Insightful Science; https://www.snapgene.com).

### Simulation of the immune system

Immunological simulations of the multi-epitope vaccine were performed using the C-IMMSIM server (https://kraken.iac.rm.cnr.it/C-IMMSIM/) to evaluate the immunogenic properties of chimeric peptides and their actual immune response profiles (89). This platform functions as an agent-based simulator of the immune system and serves as a versatile tool that applies position-specific scoring matrices (PSSM) along with machine learning techniques to predict immune epitopes and their interactions.

Simultaneously, the simulation models three distinct anatomical regions in mammals: the thymus, bone marrow, and tertiary lymphoid organs (90). To mimic the immune response, the vaccine was delivered in three doses of 1000 molecules each, spaced four weeks apart. The random seed, simulation volume, and simulation step parameters were set to 12,345, 10 L, and 1050, respectively. Previous studies indicate that the minimum interval between two injections should be four weeks. The three doses corresponded to time steps of 1, 84, and 168, each representing roughly 8 hours of real time (91). Administering vaccine doses at defined intervals is recommended to assess the effects of repeated *Helicobacter pylori* exposure on immune responses.

## Results

### Prediction of antigenicity and alignment of protein sequences

According to the antigenicity screening of the amino acid sequences of four structural proteins (Outer membrane protein and extracellular protein) from *Helicobacter pylori* (strain ATCC 700392 / 26695) pan proteome, the SabA protein had the most excellent antigenic value, followed by proteins SabB, VacA, and BabB in that order. It was also determined that these proteins could serve as desirable antigens (**Table 1 and Supplementary Table S3**). Hence, these proteins were selected to predict T and B-cell epitopes and vaccine development. After examining the results of sequence alignment, protective and non-protective regions of these structural proteins in the desired strains were identified by the identity of the amino acid sequences (Figure 2).

**Figure 2.**
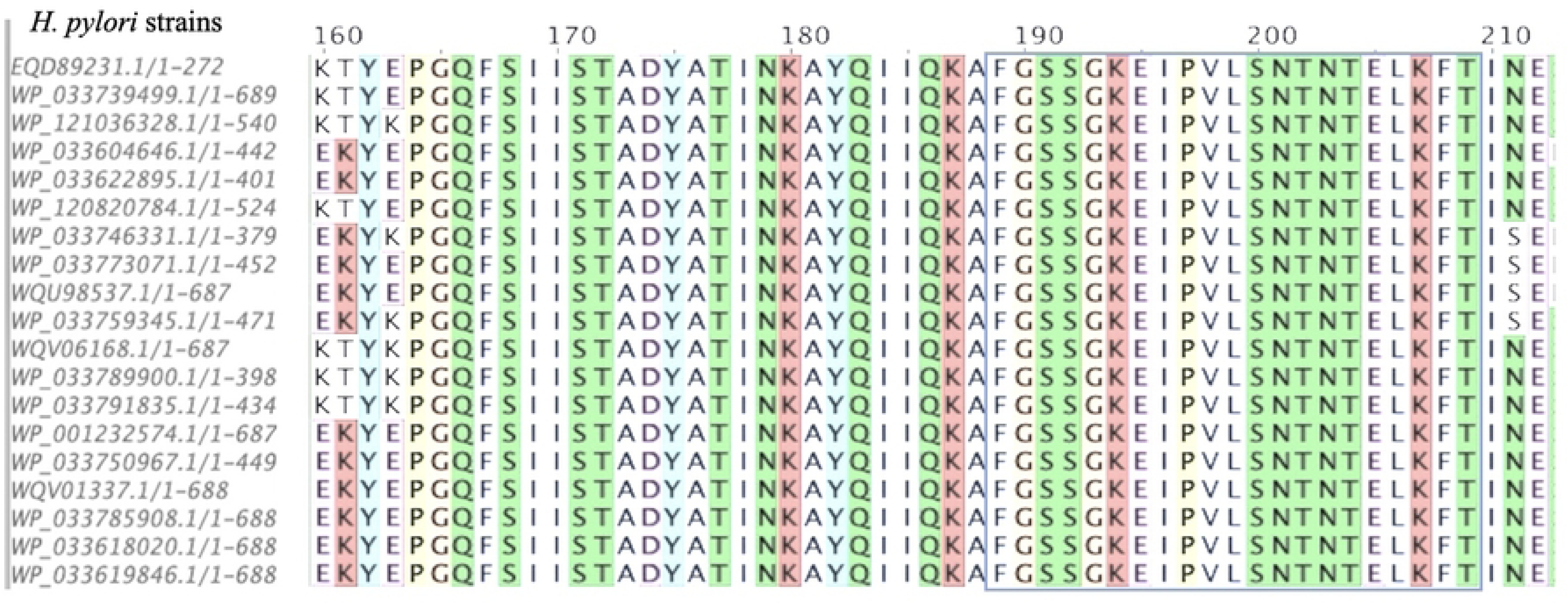
Multiple sequence alignment (MSA) for structural proteins of 19 H. pylori strains using ClustalOmega. One predicted epitope was highlighted with a blue box line in the conserved region.

**Table 1.** VaxiJen and ANTIGENpro antigenicity scores of the selected proteins and corresponding threshold values.

|  | (BabB) | (SabB/) | (SabA) | (VacA) | threshold |
| --- | --- | --- | --- | --- | --- |
| Vaxijen | 0.4938 | 0.4591 | 0.5685 | 0.5709 | 0.4 |
| AntigenPro | 0.843338 | 0.909562 | 0.952467 | 0.782333 | 0.5 |

To enhance the efficacy of the designed vaccine against both wild-type and mutant strains of *Helicobacter pylori*, our vaccine design strategy was based on the analysis of the bacterial pan-proteome. By examining the complete set of proteins across multiple strains, we were able to identify and select epitopes that are recognized by both predictive tools as stimulating T and B lymphocytes, while also being located within non-mutated and fully conserved regions of the bacteria.

This pan-proteome-based approach ensures that the vaccine targets regions less prone to mutations, increasing the likelihood of broad protection across diverse bacterial strains. Nonetheless, it is important to note that the immune response is complex and may involve additional factors beyond predicted epitope conservancy. Therefore, the immunogenic potential of selected epitopes and their ability to confer complete protection should be confirmed through experimental validation (92).

### T cell epitope prediction: CTL and HTL

The proteasome breaks down pathogen or vaccine peptides in the cytosol by unfolding the protein and cleaving it into peptide pieces. The protein transporter TAP carries them to the endoplasmic reticulum. The MHC I groove is only securely bound by peptides of the appropriate size (100). Proteasomal cleavage, TAP transport efficiency, and MHC class I binding are all evaluated using the NetCTL 1.2 server. The chance of peptide location in the MHC-I groove increases with higher predicted values for these indicators (101). NetCTL1.2 estimated 288 epitopes for BabB, 652 for SabB, 264 for SabA, and 292 for VacA protein.

These epitopes demonstrated strong affinity for binding to several of the seven MHC class I supertypes’ alleles. All of these epitopes were included in the IEDB results as selected items by taking into account all of the MHC alleles for these structural proteins that were available on this server. Conversely, 238 epitopes for BabB, 422 for SabB, 258 for SabA, and 308 for VacA were predicted by the NetMHC II pan 3.2 server, which was utilized to predict HTL epitopes. The anticipated epitopes across both servers were somewhat confirmed when it was discovered that IEDB also recommended these same epitopes as top epitopes for the alleles in this server. Among these anticipated epitopes for helper T lymphocytes (HTLs) and cytotoxic T lymphocytes (CTLs), the best ones were selected based on antigenicity, allergenicity, toxicity, and most importantly, 100% conservation in all *Helicobacter pylori* strains.

. These criteria led to the selection of 29 CTL epitopes of BabB, 36 of SabB, 41 of SabA, and 39 of VacA. Furthermore, screening was done for 23 HTL epitopes of BaB, 31 of SabB, 38 of SabA, and 29 of VacA. Eight epitopes in pr1, three in pr2, five in pr3, and five in pr4 were found to match the required criteria. **Table 2**, **Table 3** list the T-cell epitopes utilized in the final vaccine formulation.

**Table 2.**
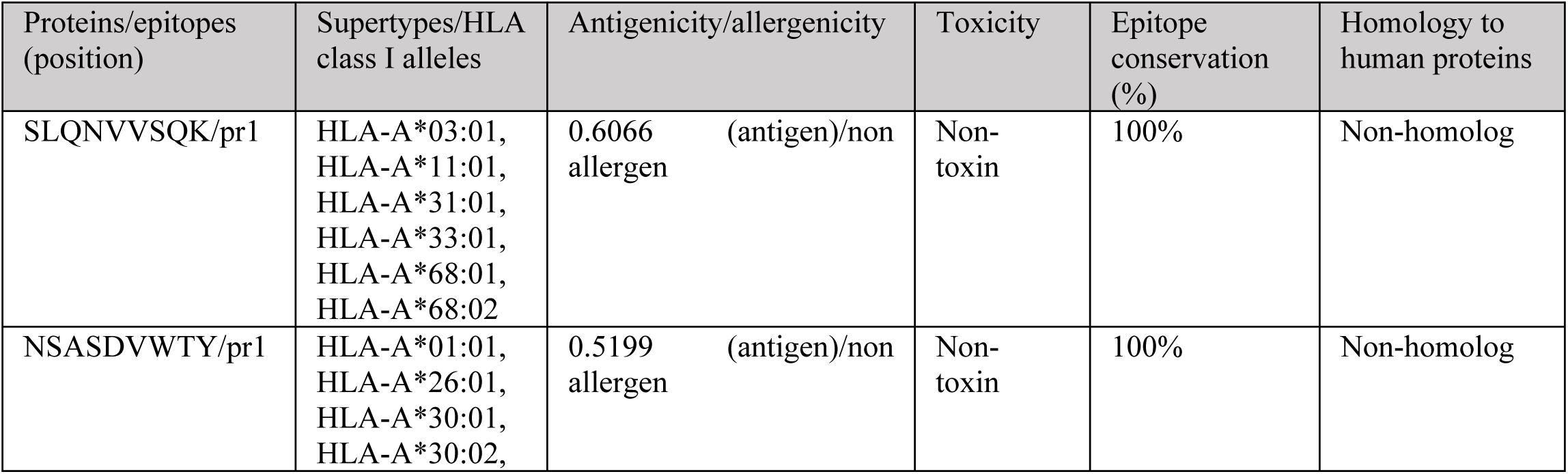
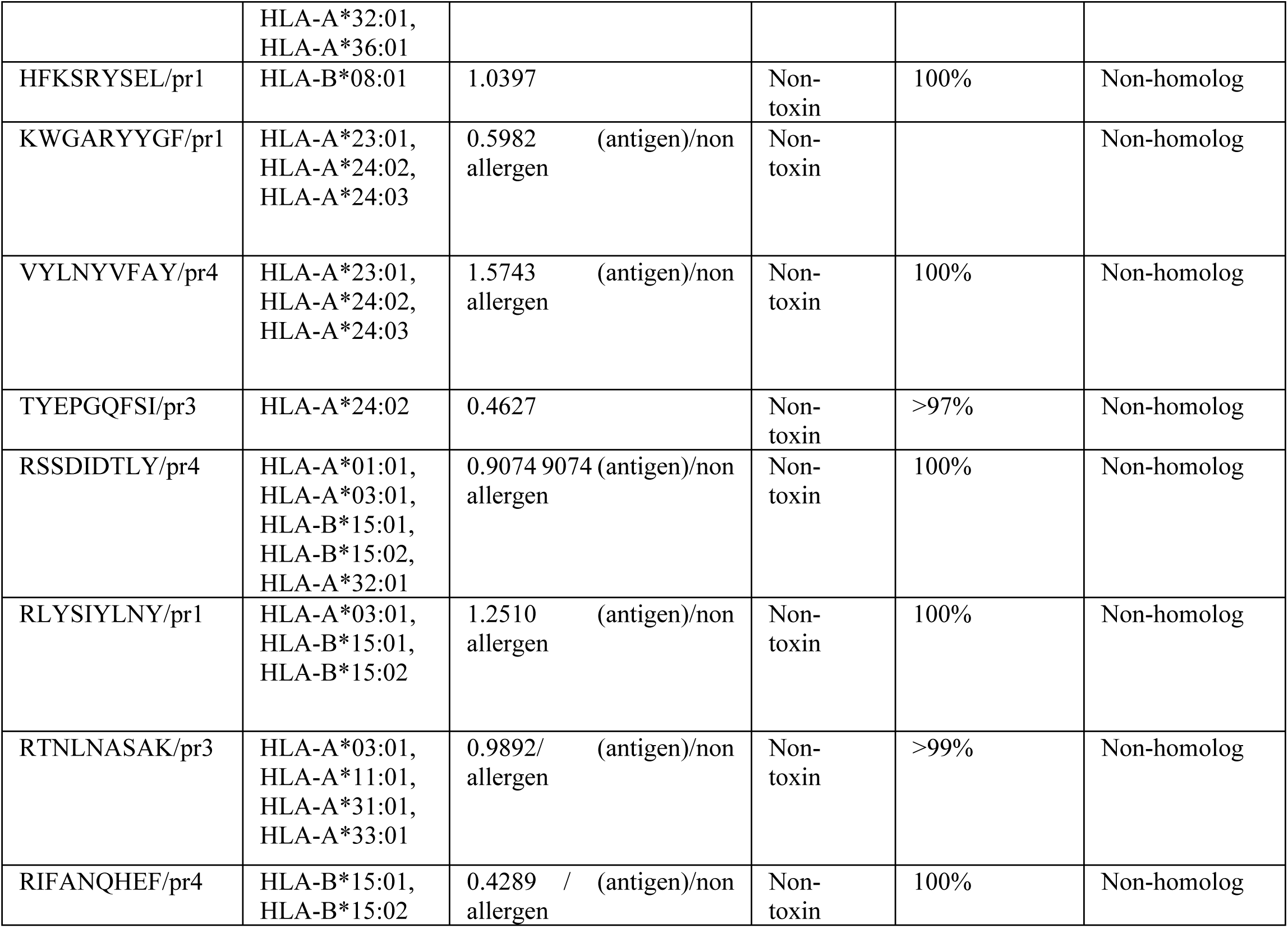
The selected CTL epitopes for the final vaccine construction are provided by the NetCTL and IEDB server.

**Table 3.**
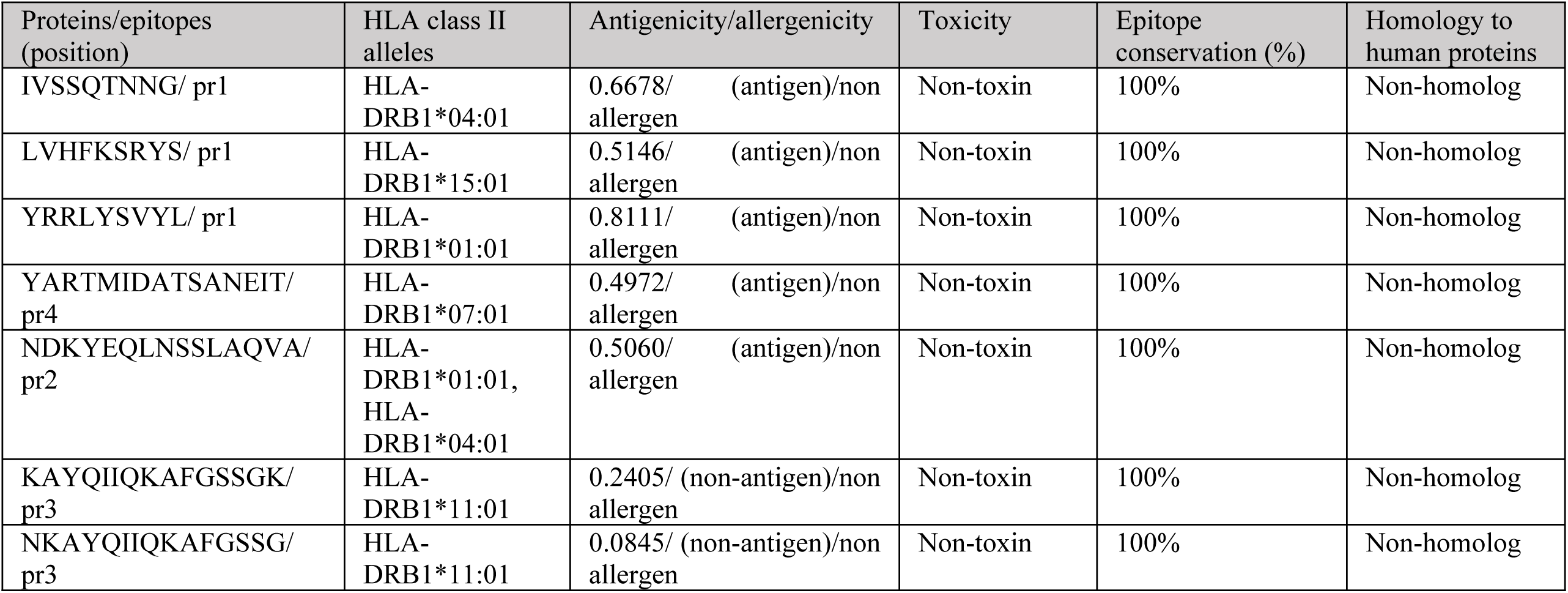
The selected HTL epitopes for the final vaccine construction are provided by the NetMHC II pan 3.2 and IEDB server.

### Linear and conformational B cell epitope prediction

A total of 27 epitopes were predicted for BabB, 14 for SabB, 12 for SabA, and 6 for VacA after closely analyzing the outcomes of five linear B-cell (LBL) epitope prediction servers and taking into account the commonalities among their outputs. Twelve linear B-cell epitopes of BabA, three of SabB, three of SabA, and two of VacA were also acquired. A number of factors, including binding score, antigenicity, allergenicity, toxicity, flexibility, hydrophilicity, surface accessibility, and complete conservation, were taken into consideration when choosing the epitopes **(Table 4**). To maximize immunogenicity and coverage, a selection of these epitopes was selected for the final vaccine design. In particular: two SabB epitopes, one SabA epitope, and one VacA epitope. The final vaccination sequence included these chosen epitopes. Six B-cell conformational epitopes were also found from the improved Three-dimensional (3D) model based on the ElliPro server results. **Table 5** displays the amino acid residues, sequence position, number of residues, and their scores. The 3D model of these epitopes is shown graphically in Supplementary Fig. S1.

**Table 4.**
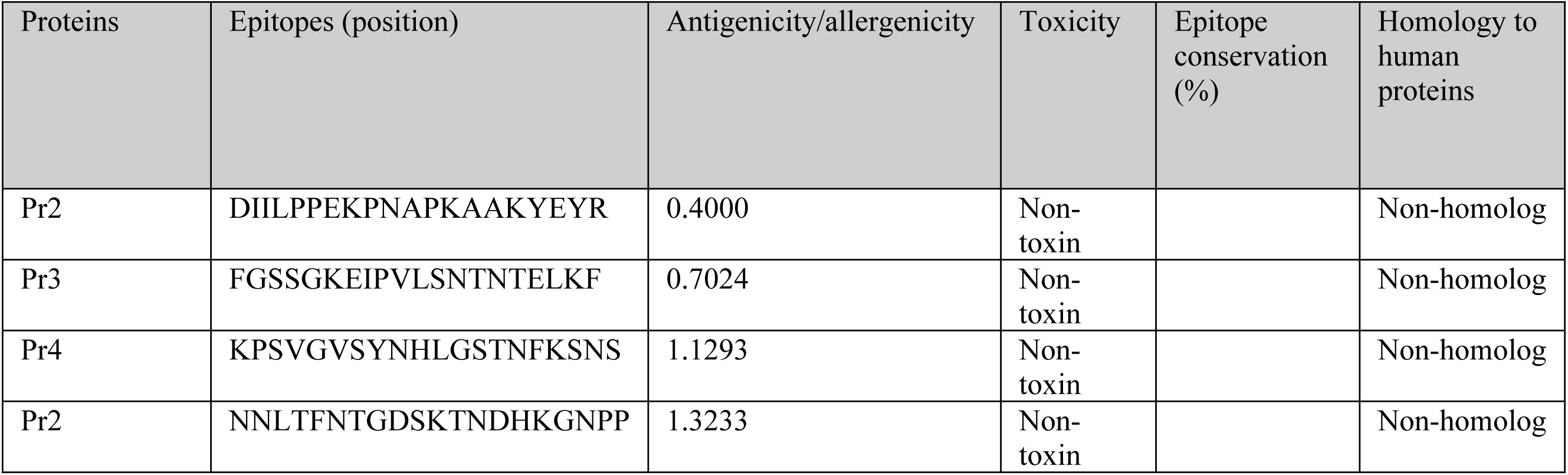
Details of linear/continuous B-cell (LBL) epitopes in the vaccine construct.

**Table 5.** List of conformational/discontinuous B-cell epitopes predicted over final vaccine construct.

| No. | Residues | Number of residues | Score |
| --- | --- | --- | --- |
| 1 | A:P395, A:N396, A:A397, A:P398, A:K399, A:A400, A:K430, A:K431, A:P432, A:S433, A:V434, A:G435, A:V436, A:S437, A:Y438, A:N439, A:H440, A:L441, A:G442, A:S443, A:T444, A:N445, A:F446, A:K447, A:S448, A:N449, A:S450, A:K451, A:K452, A:N453, A:N454, A:L455, A:T456, A:F457, A:N458, A:T459, A:G460, A:D461, A:S462, A:K463, A:T464, A:N465, A:D466, A:H467, A:K468, A:G469, A:N470, A:P471, A:P472, A:G473, A:G474, A:G475, A:G476, A:S477, A:H478, A:E479, A:H480, A:E481, A:H482 | 59 | 0.824 |
| 2 | A:H34, A:N35, A:T36, A:Q37, A:I38, A:Y39, A:T40, A:L41, A:N42, A:D43, A:K44, A:I45, A:F46, A:S47, A:Y48, A:T49, A:E50, A:S51, A:L52, A:A53, A:G54, A:K55, A:R56, A:E57, A:M58, A:A59, A:I60, A:I61, A:T62, A:F63, A:K64, A:N65, A:G66, A:A67, A:I68, A:F69, A:Q70, A:V71, A:E72, A:V73, A:P74, A:G75, A:S76, A:Q77, A:H78, A:I79, A:D80, A:Q82, A:K83, A:I86, A:E87, A:K90, A:D91, A:T92, A:L93, A:R94, A:I95, A:A96, A:Y97, A:L98, A:T99, A:E100, A:A101, A:K102, A:V103, A:E104, A:K105, A:L106, A:V108, A:W109, A:N110, A:N111, A:K112, A:T113, A:P114, A:H115, A:A116, A:I117, A:M122, A:A123, A:N124, A:E125, A:A126, A:A127, A:A128, A:K129, A:A130, A:F132, A:V133, A:A134, A:W136, A:T137, A:L138, A:A140, A:A141, A:A142, A:A143, A:A144, A:Y145, A:S146, A:L147, A:Q148, A:N149, A:V150, A:V151, A:S152, A:Q153, A:A155, A:A156, A:Y157, A:N158, A:S159, A:R174, A:Y175 | 114 | 0.736 |
| 3 | A:L299, A:Y300, A:M314, A:I315, A:D316, A:A317, A:T318, A:S319, A:A320, A:N321, A:E322, A:I323, A:T324, A:G325, A:P326, A:G327, A:P328, A:G329, A:N330, A:D331, A:K332, A:F359, A:G360, A:S361, A:S362, A:G363, A:K364, A:G365, A:P366, A:G367, A:P368, A:G369, A:N370 | 33 | 0.652 |
| 4 | A:F262, A:G263, A:P264, A:G265, A:I268, A:V269, A:S270, A:S271, A:Q272, A:T273, A:N274, A:N275, A:G276, A:G277, A:P278, A:G279, A:P280, A:G281, A:L282, A:G295, A:R298 | 21 | 0.636 |
| 5 | A:M1, A:I2, A:K3 | 3 | 0.623 |
| 6 | A:V418, A:L419, A:S420, A:N421, A:T422, A:N423, A:T424, A:E425, A:L426, A:K427 | 10 | 0.56 |

### Analysis of human population coverage around the world, protection assessment, and auto-immunity identification

The human population coverage analysis demonstrated that the 17 selected T-cell epitopes in this study cover approximately 92.13% of the global population. Regional coverage percentages are detailed in **Table 6 and Supplementary Fig. S2**. This indicates that the designed multi-epitope vaccine is broadly applicable for combating Helicobacter pylori across most regions. Notably, North America showed the highest coverage at 95.15%, whereas Central America had the lowest at 22.09%. The reduced coverage in Central America is attributed to the region’s complex genetic admixture from multiple migration waves, resulting in diverse ancestral population proportions. Population genetics studies reveal significant variation in HLA profiles correlating with these admixture levels, which likely affects epitope recognition (93). Because Central American populations exhibit substantial genetic diversity, a peptide vaccine based on specific epitopes not prevalent in their genetic makeup may result in decreased epitope coverage. Conservation analysis using the IEDB tool confirmed that the selected T- and B-cell epitopes are highly conserved among different species, particularly in regions without alterations in the studied structural protein sequences. Furthermore, all selected epitopes showed no homology with the human proteome, supporting that the multi-epitope vaccine is likely to elicit a safe antigenic response without cross-reactivity to human proteins, minimizing the risk of autoimmune reactions.

**Table 6.** Population coverage of the selected epitope included in the vaccine construct. a Coverage of population on projected. b Population recognized by HLA combinations/epitope hits on the average number.

| Population/area | Class combined |  |  |
| --- | --- | --- | --- |
|  | Coverage<sup>a</sup> | Average hit<sup>b</sup> | pc90<sup>c</sup> |
| World | 92.13 | 3.28 | 1.11 |
| East Asia | 83.1 | 3.44 | 0.59 |
| Northeast Asia | 65.35 | 2.19 | 0.29 |
| South Asia | 94.30 | 3.62 | 1.45 |
| Southeast Asia | 84.26 | 2.57 | 0.54 |
| Southwest Asia | 84.26% | 2.57 | 0.64 |
| Europe | 78.35 | 2.25 | 0.59 |
| East Africa | 85.15% | 2.39 | 0.67 |
| West Africa | 88.92% | 2.72 | 0.9 |
| Central Africa | 77.35% | 2.1 | 0.44 |
| North Africa | 91.5% | 2.93 | 1.11 |
| South Africa | 86.82% | 2.32 | 0.76 |
| North America | 95.15% | 3.98 | 1.62 |
| Central America | 22.09% | 0.28 | 0.13 |
| South America | 82.13% | 2.49 | 0.56 |
| Average | 81.07 | 2.57 | 0.78 |
| Standard deviation | 16.57 | 1.00 | 0.39 |

### Creating the final multi-epitope vaccine

The epitopes CTL, HTL, LBL, Pan HLA-DR reactive epitope (PADRE), adjuvant (cholera toxin B subunit/CTB), linkers (EAAAK, AAY, GPGPG, KK, GGGGS), and the H5E tag comprise the linear vaccine’s structure. The main layout of this structure is shown in Figure 3.

**Figure 3.**
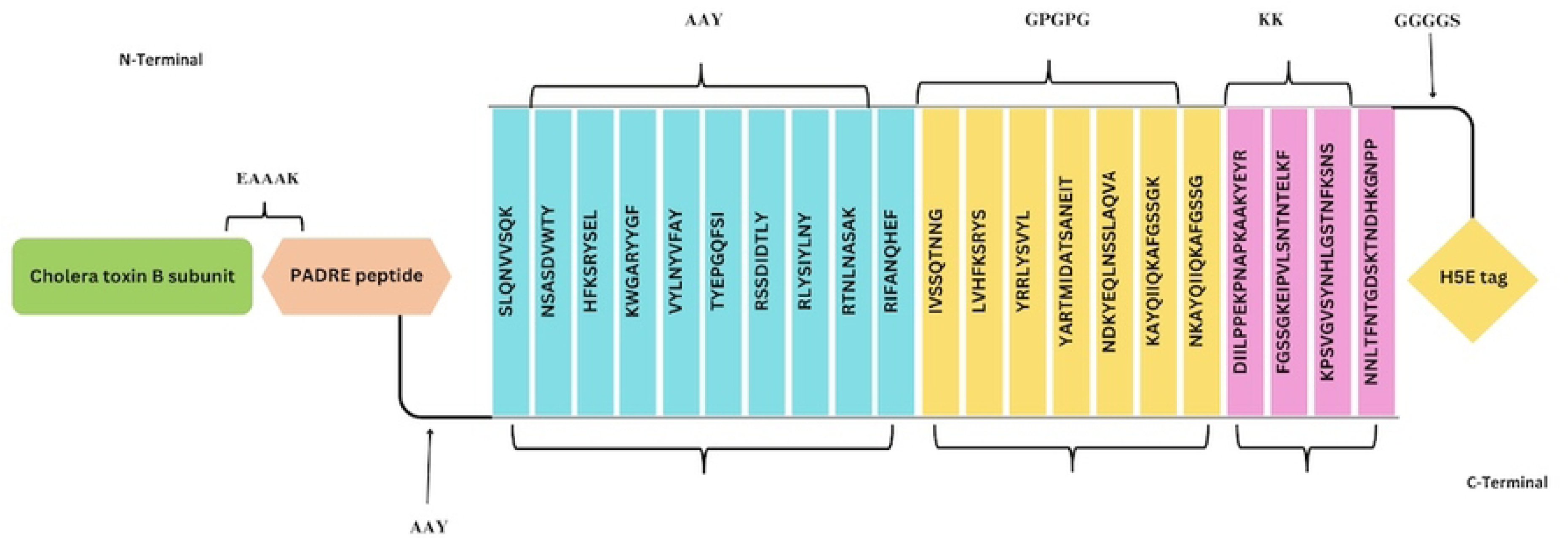
Schematic Presentation of the final multi-epitope vaccine construct.

### Assessment of the vaccine’s antigenicity, allergenicity, toxicity, solubility, and physicochemical characteristics

Assessment of the vaccine’s antigenicity, allergenicity, toxicity, solubility, and physicochemical characteristics The final sequence of the vaccine was evaluated for antigenicity, allergenicity, toxicity, and solubility, and it satisfied every requirement. The resulting vaccination has a high antigenicity, as confirmed by VaxiJen v2.0 servers (94) with a score of 0.7359 and ANTIGENpro (95) with a value of 0.914034. The final vaccination is non-allergenic and does not cause allergic reactions, according to AllergenFP 1.0 and AllerTOP 2.0 servers (52,53) demonstrated that the finished vaccination does not cause allergic reactions and is non-allergenic. The complete final vaccination sequence, including the adjuvant sequence, all epitopes, linkers, PADRE, and H5E tag, does not contain any toxin portion, according to the results of the SVM prediction mode in the ToxinPred service (54). The vaccine has acceptable solubility, according to the values predicted by Protein-sol server (109) (0.465) and SolPro server (55) (0.655634) and Protein-sol server (56) (0.465) also show that the vaccine has good solubility (Table 7). Because it affects the distribution, stability, efficacy, formulation, administration, adsorption, manufacturing process, storage conditions, bioavailability of the vaccine components, and overall success of vaccination strategies, solubility is a crucial factor in vaccine development (96).

**Table 7.** Antigenicity, allergenicity, solubility, toxicity, and physicochemical properties prediction results for final vaccine construction.

|  |  |  |
| --- | --- | --- |
| **Antigenicity** | Vaxijen | 0.7359 |
|  | AntigenPro | 0.914034 |
| **Allergenicity** | AllerTOP | Non-allergen |
|  | AllergenFP | Non-allergen |

| Toxicity | Non-toxin |  |
| --- | --- | --- |
| solubility | SolPro | 0.655634 |
|  | Protein-sol | 0.465 |
| Physicochemical properties | Mol. weight (kDa) | 53130.81 |
|  | Theoretical pI | 9.53 |
|  | Half-life (in vitro) | 30 hours |
|  | Half-life (in vivo) | >20 hours (Yeast)<br>>10 hours (*Escherichia coli*) |
|  | Instability index | 19.63(stable) |
|  | Aliphatic index | 66.21 |
|  | Hydropathicity (GRAVY) | -0.484 |

The final structure comprises 486 residues and a molecular weight of 53130.81 kDa, according to the findings of the assessment of the vaccine’s physicochemical characteristics. This structure is crucial, as evidenced by its theoretical isoelectric point (pI) of 9.53. Furthermore, the vaccine’s estimated half-lives in mammalian reticulocytes (in vitro), yeast (in vivo), and E. coli (in vivo) are 30 hours, 20 hours, and 10 hours, respectively, indicating the vaccine’s stability in various phases. A vaccine’s half-life varies depending on the organism, which shows how stable and efficient it is in various biological settings.

The dosing schedule and practicality may be impacted if a shorter half-life necessitates more frequent administration. extended half-lives, however, can provide protection for extended periods of time, but they may also raise safety issues or necessitate quick adaption to new strains. The nature of the vaccine, the target pathogen, and practical administration factors all affect the ideal half-life (97). According to the forecast, the vaccine’s structure has a high degree of stability to trigger an immunological response, with an instability score of 19.63 (< 40). The vaccine’s aliphatic index is 66.21, indicating that it is thermostable. A protein becomes heat resistant as its aliphatic index rises.

The overall average hydrophilicity (GRAVY) was-0.484. Lower GRAVY values imply better solubility, indicating that the vaccine candidate is hydrophilic, enhancing interaction with water and blood and enabling the identification of the Ψ target easier (98). The vaccine structure satisfies the requirements for vaccine formulation, according to all of the physicochemical properties study results (**Table 7 and Supplementary Material SM1**). A difficulty with expression during vaccine manufacture is not anticipated because the planned vaccine lacks any transmembrane helices. Furthermore, the vaccine construct’s signal peptides promote strong humoral immune responses against bacterial pathogens and improve B cell identification of antigens by facilitating protein production and extracellular localization. (28) (**Supplementary Figs. S3 and S4**).

### The vaccine construct’s secondary structure

Using PSI-blast-based secondary structure prediction (PSIPRED) (99), the secondary structure of the vaccine was predicted to be composed of 41% helix, 14.40% strand, and 44.40% coil. The Self-Optimized Prediction Method with Alignment (SOPMA) (60) tool also supplied the secondary structure, with default parameters of 10.91% alpha-helix, 10% extended strand, and 79.01% random coil. The secondary structural features are displayed graphically in Figure 4 and Supplementary Figure S6. As can be seen from the picture, the high percentage of random coil suggests that epitopes are present in various parts of the Ψ construct (100).

**Figure 4.**
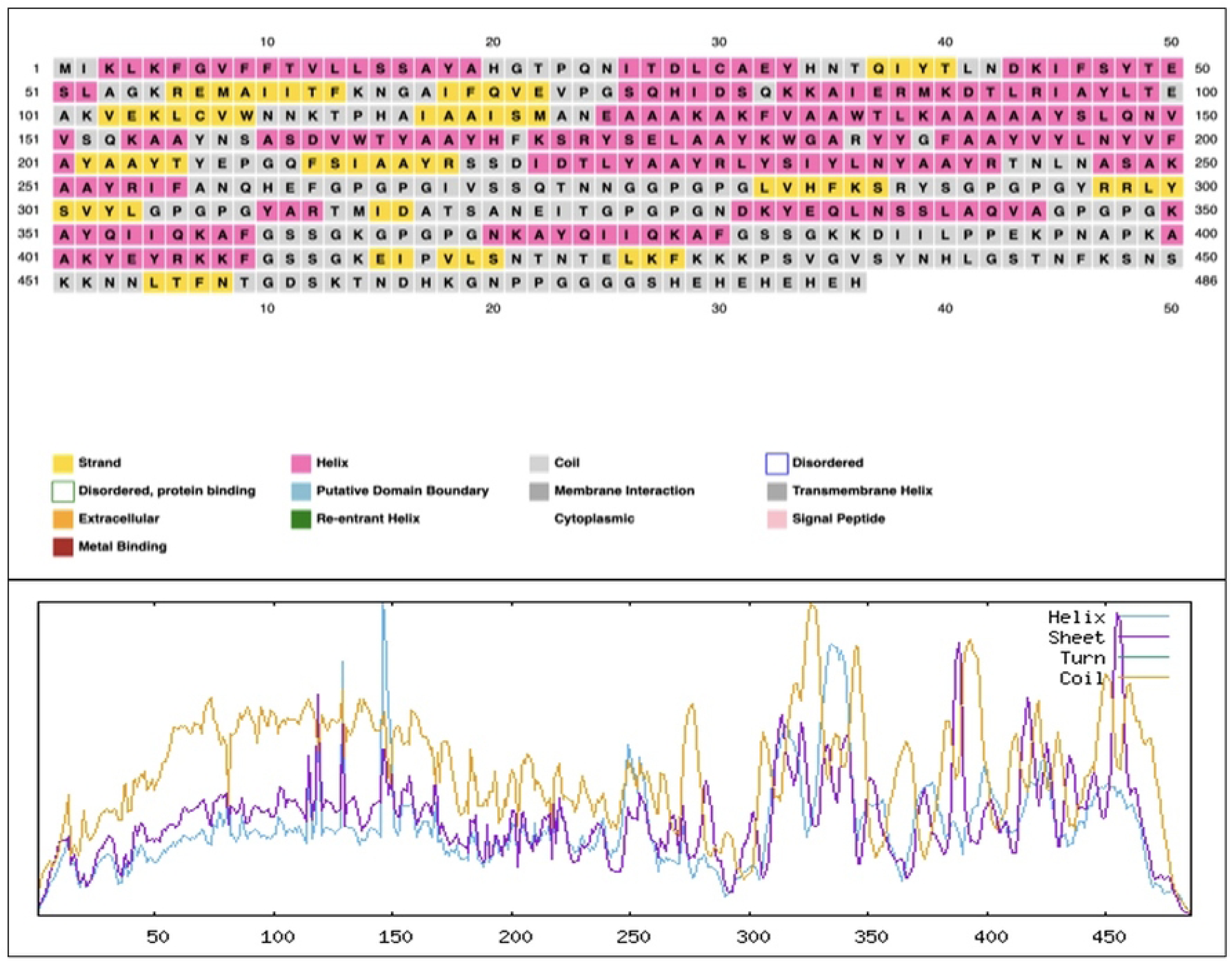
Secondary structure prediction of the vaccine, performed using the SOPMA server.

### Modeling, improving, and verifying the multi-epitope vaccine’s three-dimensional structure

After creating the 3D vaccine model using the Robetta server (62), one model was selected as the best prediction model, and based on this model, the 3D structure of the final vaccine was made (Supplementary Figure S7). Later than refining the 3D structural model obtained from the modeling stage by GalaxyRefne server (101), model 1 was selected as the best final vaccine model based on various parameters, including GDT-HA (0.9835), RMSD (0.294), MolProbity (1.795), Clash score (10.2), weak rotamers (0.0), and Rama favored (96.1) among other refined models (**Figure 5a** and Supplementary Table S4). The Z-score for the vaccine structure in the PROSA diagram was −7.8. This score is close to the range of native proteins of similar size, indicating the lowest error rate and accuracy of the simulation and the overall reliability of the predicted model (**Figure 5b**, c). In addition, the quality factor of 89.565 obtained using the ERRAT server indicates the optimal quality of the protein model (102) (**Figure 5d**). Grouping of amino acids based on phi and psi angles by Ramachandran plot (77,103) using PROCHECK analysis in the PDBsum server for the refined structure, revealed that 91.5% of the residues were classified in the most favorable, 7.2% additional allowed, 0.0% generously allowed, and 1.2% disallowed region (**Figure 5e**).

**Figure 5.**
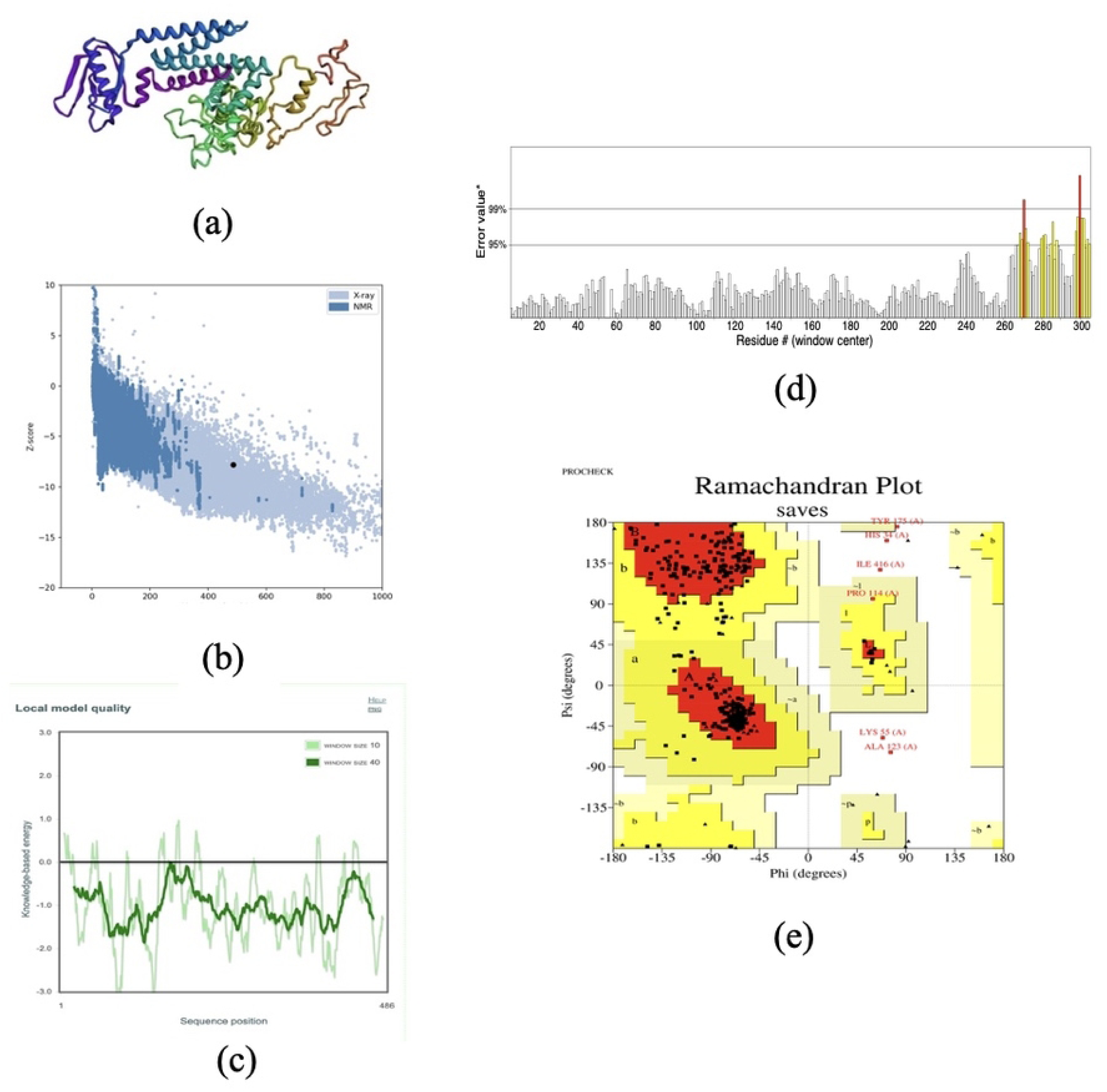
(a) The three-dimensional refined vaccine model is visualized to represent the helical,sheet, and loop regions. (b) ProSA validation of predicted structure with Z-score of *9.78. (c) plots the residues scores to check the local model quality. (d) ERRAT plot illustrating error values (%) along the protein sequence using a sliding window approach, with regions exceeding 95% and 99% error thresholds highlighted in yellow and red, respectively. (e) Analysis of the Ramachandran plot utilizing the PROCHECK server showed 94.6%, 3.5%, 0.4%, and 1.5% residues laying in favored, additional allowed, allowed, and disallowed regions, respectively.

According to the results, the total residuals in the desired area were in the ideal value range, i.e., more than 90%, which confirms the reliability of this model. The structural quality of the refined vaccine model was further validated using the Verify3D server. The results showed that 86.01% of the residues exhibited an averaged 3D–1D score ≥ 0.1, exceeding the acceptable threshold of 80%. This indicates good compatibility between the predicted three-dimensional structure and its amino acid sequence, supporting the reliability of the modeled vaccine structure Supplementary Figure S8.

### Molecular docking of the vaccine construct with TLR4, MHC-I, and II receptors and binding affinity evaluation

Protein–protein docking between the refined 3D model of the final vaccine construct and the immune receptors (TLR4, MHC-I, and MHC-II) was performed using the ClusPro 2.0 web server. The generated docking poses were grouped into multiple clusters based on their structural similarity (RMSD-based clustering). Each cluster represents a set of energetically favorable models, ranked according to cluster size and the weighted energy score of the cluster center. The most reliable docking solutions were selected from clusters showing the lowest energy score and the largest number of members.

Table 8 summarizes the statistical parameters and their corresponding values for each docked vaccine–receptor complex. Supplementary Figs. S12–S14 present the graphical docking results generated by the ClusPro server for each receptor–vaccine complex. The weighted energy score, cluster size, RMSD values, and other ClusPro-reported parameters were evaluated for all predicted complexes. In ClusPro, a lower weighted energy score (more negative value) indicates a more favorable and stable interaction. Additionally, the buried surface area (BSA) reflects the extent of the interface region, with higher BSA values suggesting tighter binding and reduced solvent exposure of the interacting surfaces. RMSD values between the cluster members and the cluster center were also considered, as low RMSD values indicate structurally consistent models and support the reliability of the predicted docking cluster.

**Table 8.** Binding affinities of the docked complexes of the vaccine with MHC-I, MHC-II, and TLR4, as predicted by the PRODIGY server.

| Complexes | Gibbs free energy (kcal mol$^{-1}$) | $K_d$ (M) |
| --- | --- | --- |
| Vaccine-MHC1 | -15.9 | 2.2e-12 |
| Vaccine-MHC2 | -27.1 | 1.3e-20 |
| Vaccine-TLR4 | -8.9 | 2.8e-07 |

An overview of molecular docking of the vaccine structure and receptors are shown in (**Figure 6a, Figure 7a, Figure 8a**). According to PDBsum results, there were 21residues of the vaccine and 27 residues of TLR4 (ChainA) in the complex between the vaccine and TLR4, and the interface area (Å2) for the vaccine and TLR4 was 1470 and 1459, respectively. The interacting residues of the vaccine-MHC-I complex were 35 residues for the vaccine and 41 residues for MHC-I (ChainA), and the interface region was 1858 Å2 for the vaccine and 1794 Å2 for MHC-I. Also, for the vaccine-MHC-II complex, 41 vaccine residues interacted with 53 MHC-II (ChainA) residues, and the interface region for the vaccine was 2503 Å2, while this region was 2402 Å2 for MHC-II. On the other hand, the molecular interaction established between the vaccine with TLR4, 4 salt bridges, 21 hydrogen bonds, and 185 non-bonding contacts; with MHC-I, 4 salt bridges, 23 hydrogen bonds and 239 non-bonded contacts and finally with MHC-II; there were 7 salt bridges, 37 hydrogen bonds and 379 non-bonded contacts (**Figure 6b, Figure 7b, Figure 8b**) and Supplementary Material SM 2, 3, 4). Hydrogen bonds are essential in molecular recognition because they are important in achieving a stable compound (9).

**Figure 6.**
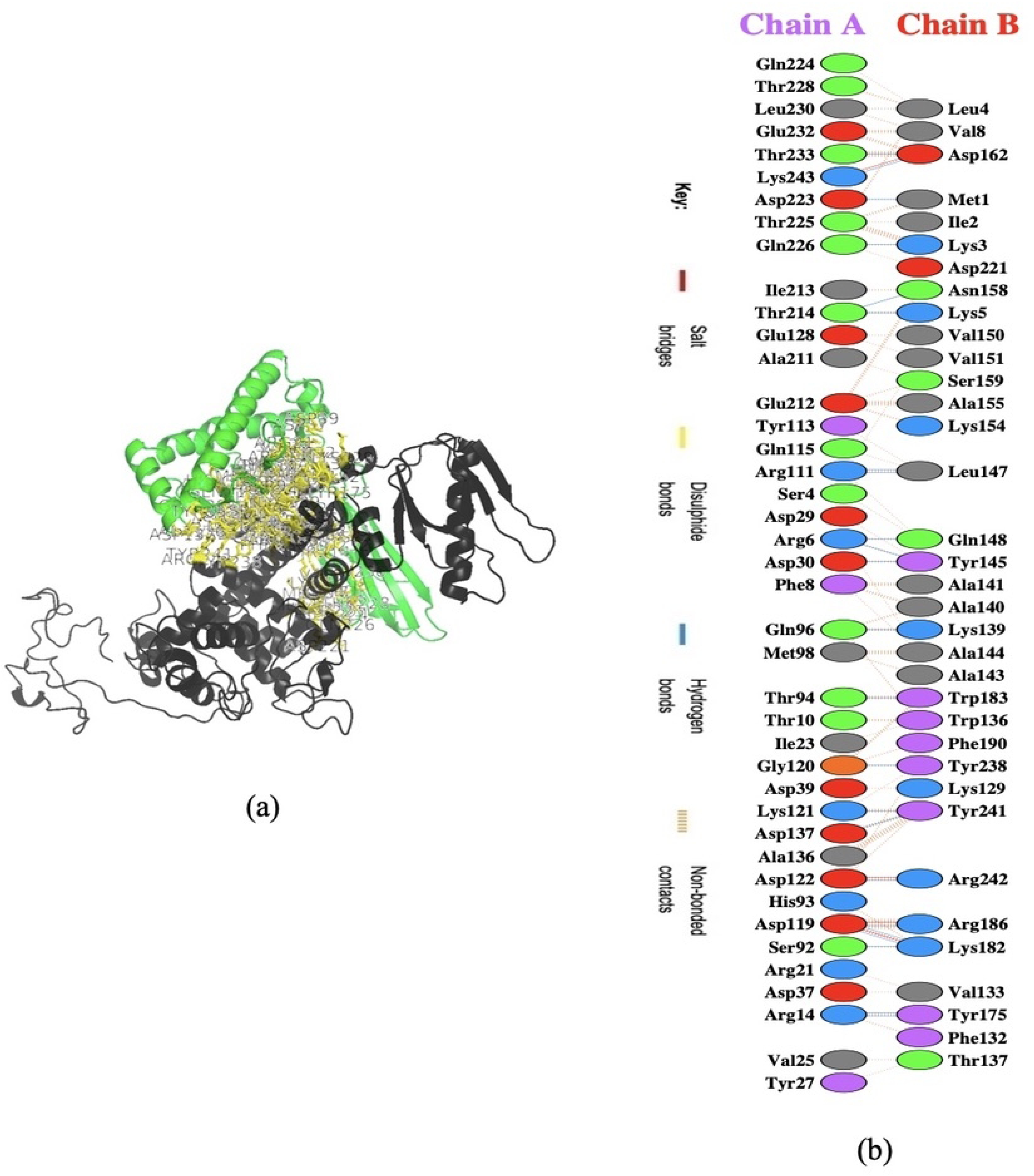
(a) Visualization of docking results for the vaccine-MHC-I complex. The vaccine construct is shown in red, while MHC-I is depicted in violet. **(b)** Map of total interacting residues and bonds between the vaccine and MHC-I protein chains.

**Figure 7.**
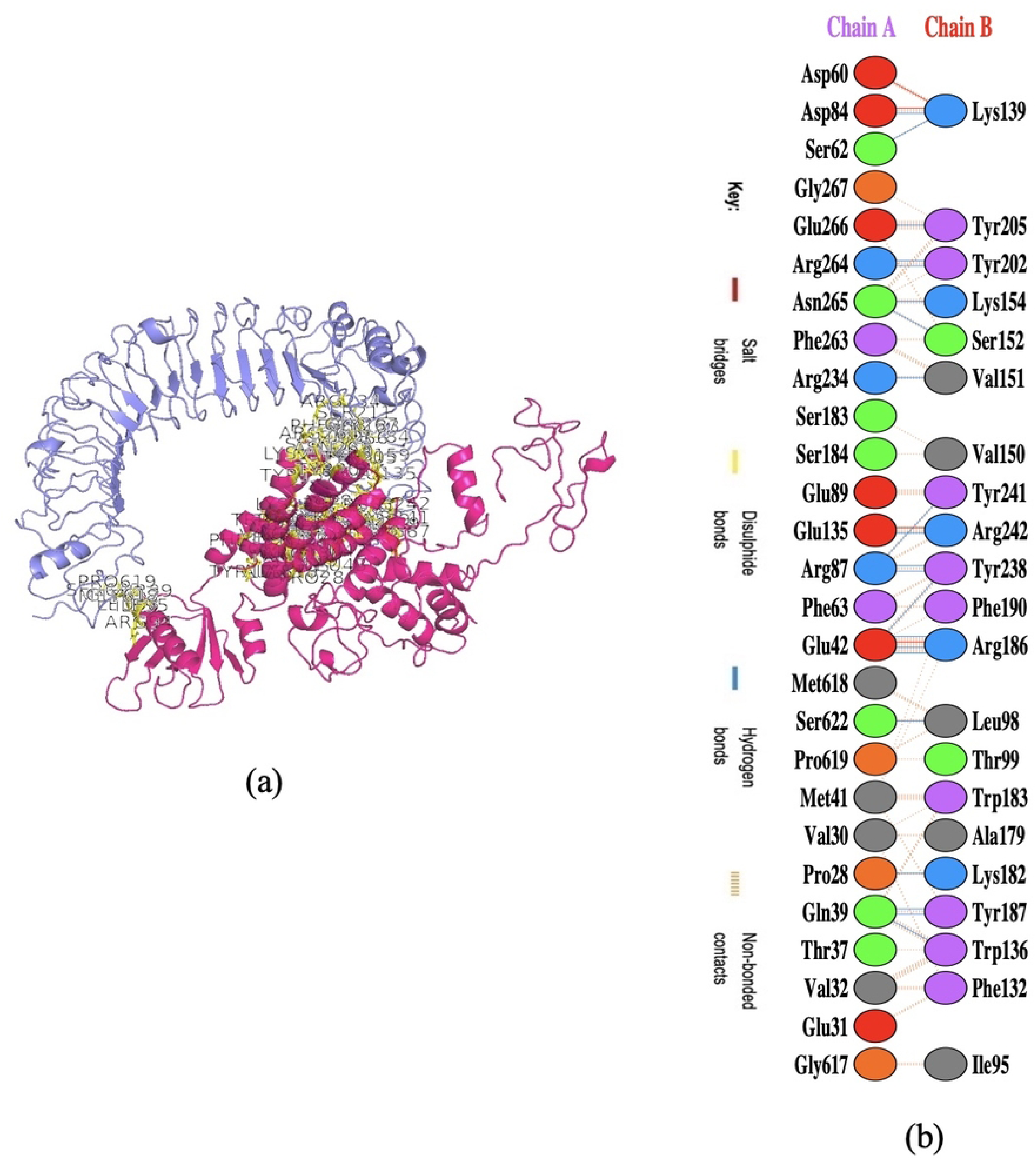
(a) Visualization of docking results for the vaccine-TLR4 complex. The vaccine construct is shown in pink, while TLR4 is depicted in orange. **(b)** Map of total interacting residues and bonds between the vaccine and TLR4 protein chains.

**Figure 8.**
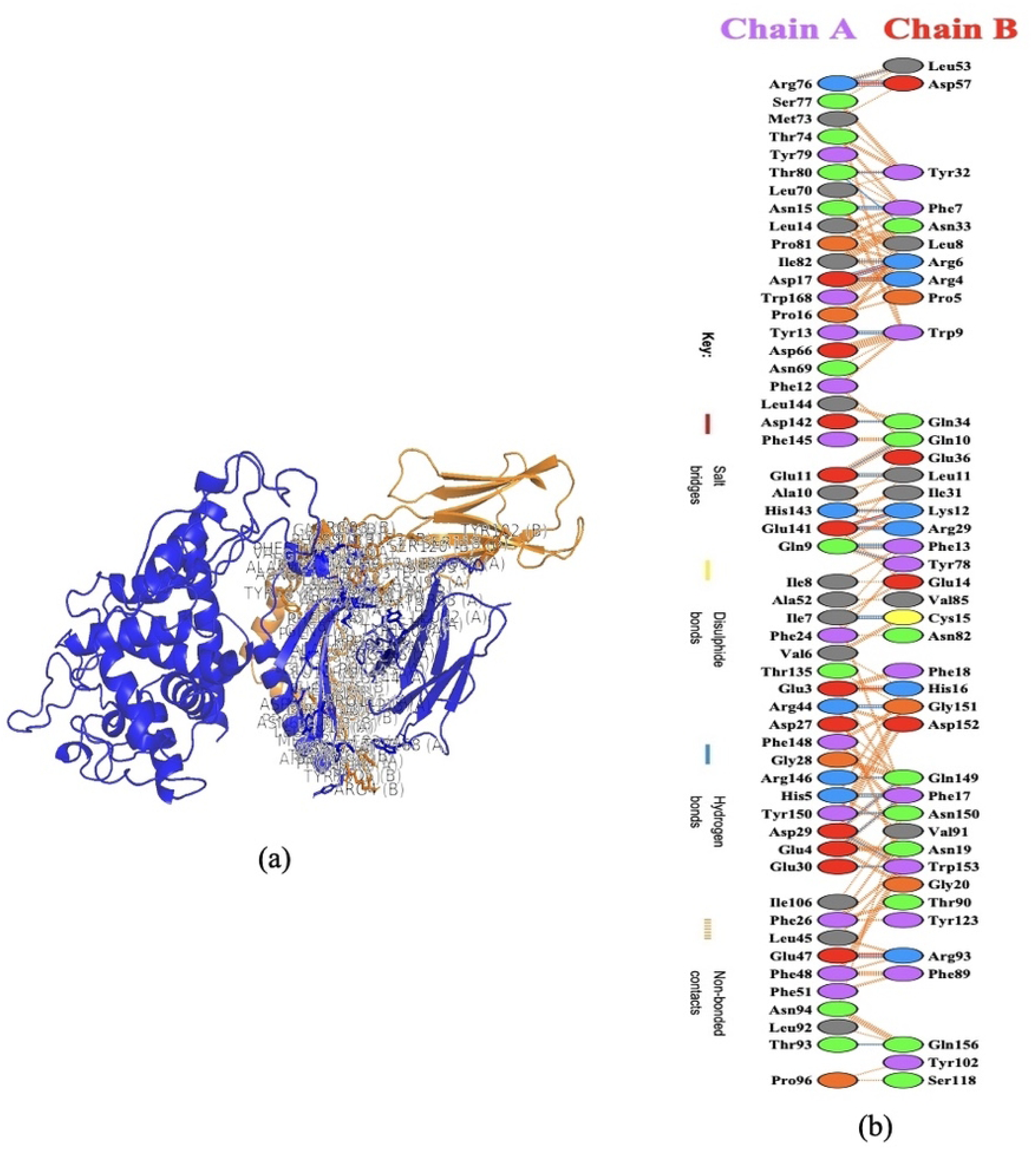
(a) Visualization of docking results for the vaccine-MHC-II complex. The vaccine construct is shown in red, while MHC-II is depicted in green. **(b)** Map of total interacting residues and bonds between the vaccine and MHC-II protein chains.

Electrostatic bonds also play an essential role in the interactions produced in complex (9). Since, in this study, most of the hydrogen bond distances between vaccine-receptor interacting residues are around 2–3 Å, there is a high interaction between them (104). Salt bridges can be essential for preserving the stability of the contact interface in protein–protein complexes. Also, they can contribute to the specificity and affinity of the interaction between two proteins and be involved in the recognition and binding of the interacting partners (105). In addition, the Ramachandran plots presented in the PDBsum results were also investigated for structural validation of the docked sets, which were also confirmatory (Supplementary Figs. S9–S11).

After binding affinity analysis using the PRODIGY (PROtein binDIng enerGY prediction) web server, ΔG values (Gibbs free energy) for TLR4-vaccine, MHC-I-vaccine, and MHC-II-vaccine complex were obtained −8.9, −15.9, −27.1 kcal mol−1 (kilocalories per mole), respectively.

All four docked complexes are energetically viable, as indicated by the negative values of ΔG.

The dissociation constant (Kd) presents binding affinity and bond strength. As binding affinity and Kd share an inverse correlation, a lower Kd signifies a heightened binding affinity. This implies that vaccine-receptor bonds are securely and tightly bound when the dissociation constant is lower (**Table 8)**. In addition to complex-level analysis, the intrinsic dynamic behavior of the vaccine construct and individual receptors was also evaluated using iMODS. The results indicated stable structural characteristics with no abnormal flexibility, supporting the suitability of the modeled structures for subsequent docking and interaction analyses (**Supplementary Figures S15–S17**).

Vaccine structural simulation using molecular dynamics and energy minimization In this study, the stability and physical motions of the vaccine structure under different Ψ conditions were estimated and evaluated using molecular dynamics simulation (MDS), a method for analyzing the atomic behavior of molecular systems (125). For the proposed vaccine, molecular dynamics modeling is used to see how it works in an actual biological system (126). Some analyses, such as energy minimization, pressure assessment, temperature, and potential energy calculations, were performed with the help of GROMACS software (127). The vaccine’s energy was minimized using the steepest descent technique, and the protein’s energy was deemed minimized when it fell below 1000 kJ·mol⁻¹.

For 2,482 steps, when the force was less than 1000 kJ·mol⁻¹, energy minimization was used. With an average potential energy of −3.26 × 10² kJ·mol⁻¹ and a total drift of −1.78 × 10◦ kJ·mol⁻¹, the system’s potential energy was determined to be −3.26 × 10¹ kJ·mol⁻¹. After 50,000 NVT steps, the average temperature was 301.8 K, with a 3.6 K temperature drift (**Figure 9A**). With a total drift of 3.6 × 10³ bar, the system’s pressure was −3.54 × 10³ bar (**Figure 9B**). With a total drift of 138.7 kg·m⁻³, the system’s computed average density was 1011.2 kg·m⁻³ (**Figure 9C**).

**Figure 9.**
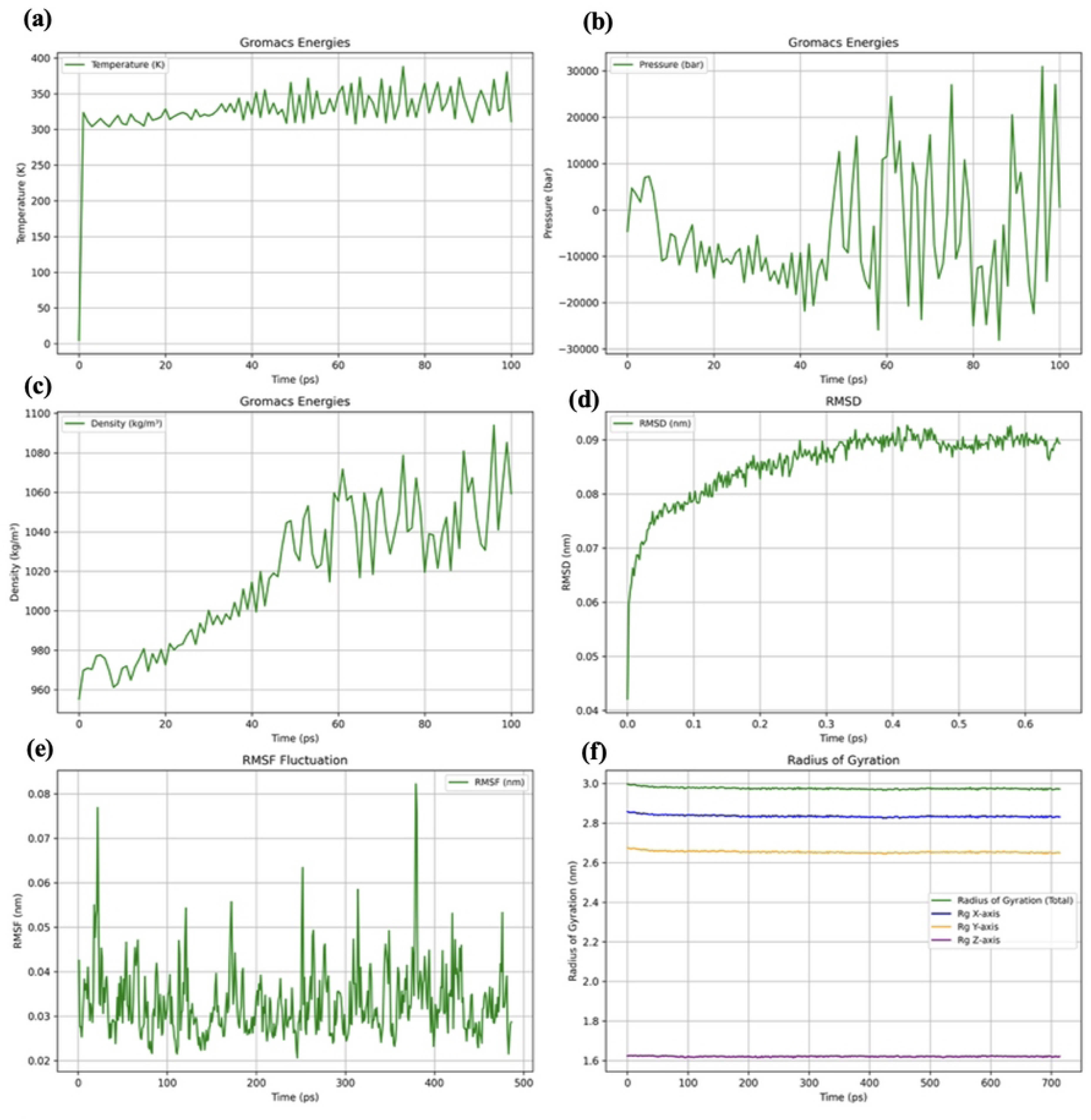
The results of molecular dynamics simulation of vaccine for analysis of structural stability. (a) Graph showing the equilibrated temperature during energy minimization. (b) Graph showing the pressure of the system during simulation. (c) Graphical presentation of density during simulation. (d) RMSD plot of the vaccine construct indicating stability. (e) RMSF plot illustrates high fluctuations, the peak-like regions with a higher degree of flexibility. (f) The Rg plot showing the vaccine construct stays compact around its axes, supporting its stability during simulation. (Data visualization and analysis performed using Python (Version 3.13.5; VS Code environment).

A trajectory analysis was carried out to verify the candidate vaccine’s stability and flexibility after a simulation period of 100 ps. The RMSD graphic communicates the stability of the vaccination over time and shows variations in its general structure. The vaccine’s RMSD fluctuated only slightly throughout the simulation, suggesting that the construct is stable (**Figure 9D**). The flexibility of the vaccine structure is shown by the RMSF plot, especially the high peaks (**Figure 9E**). Furthermore, the radius of gyration plot (Rg) (**Figure 9F**) verified the structure’s stability and compactness during the simulation.

### Codon optimization and vaccination in silico cloning

Following codon optimization (JCat) to enhance protein expression in the E. coli strain K12 as the expression host organism (106) (Supplementary Table S5). The guanine-cytosine content (GC-Content) and codon adaptation index (CAI Value) for the 1450 nucleotide optimized codon sequence were 49.10 and 0.9744, respectively (Supplementary Material SM5). Good protein expression in the host system is often indicated by a GC between 30 and 70% (98) and a CAI above 0.8, or nearly 154. This further validates the vaccine developed in this strain’s practical expression.

The cloning of the vaccine sequence within the pET-28a (+) vector to create a recombinant plasmid for creating an effective in silico cloning method is depicted in Figure 10, the result of the SnapGene software (https://www.snapgene.com/).

**Figure 10.**
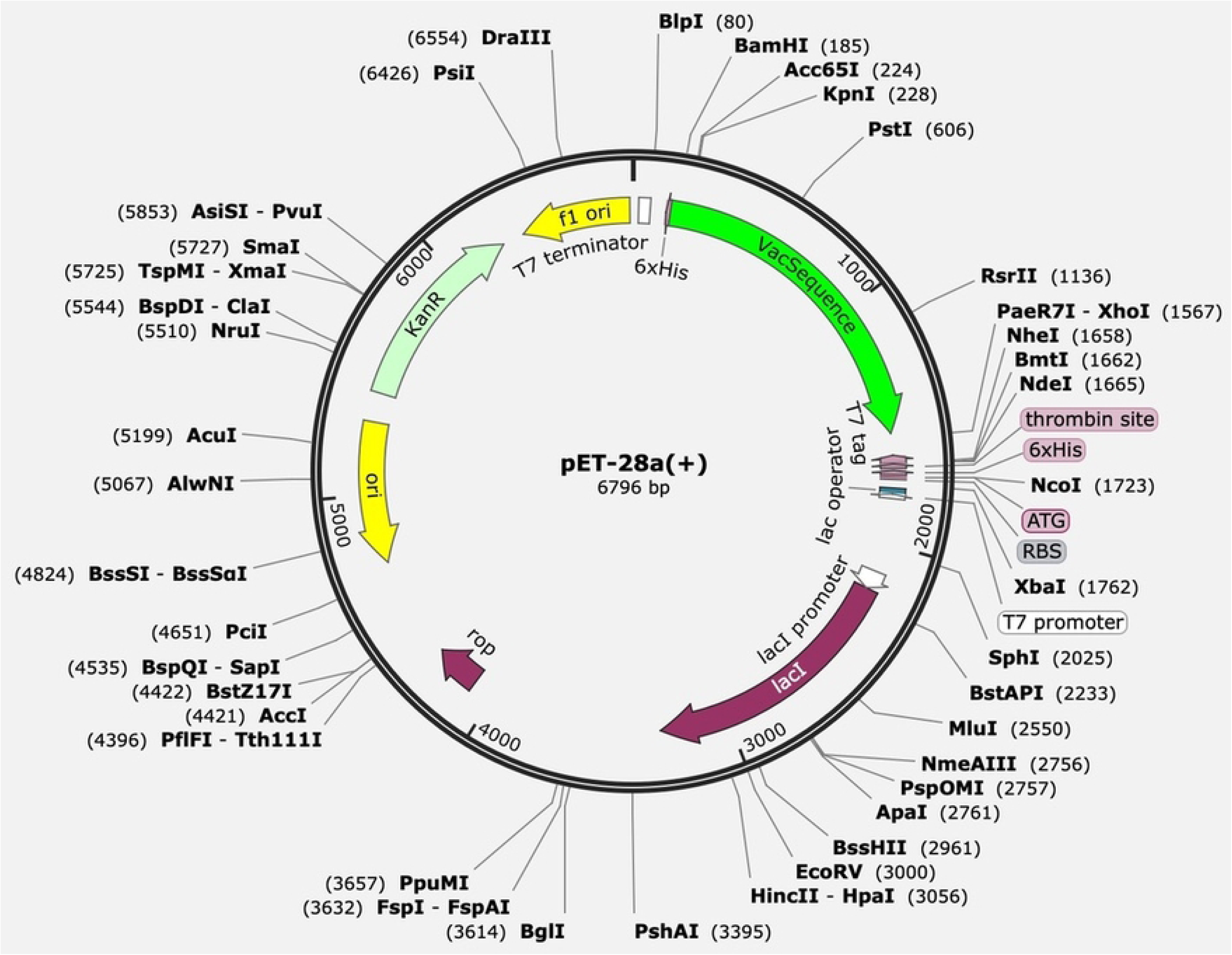
In silico restriction cloning of the designed vaccine into the pET-28a (+) expression vector. The green bar represents the codon-optimized gene of the vaccine, and the black circle represents the vector backbone.

### Immune modeling

A variety of immunoglobulins (96) may be induced by administering this vaccine with three injections, according to immunological simulation results (89) Figure. IgM levels rise in the first response, while levels of IgM + IgG, IgG1 + IgG2, IgG1, and IgG2 were considerably greater in the simulation’s second and third responses. The concentration of certain antigens with normally high immunoglobulin concentrations, such as IgG1 + IgG2, IgM, and IgG + IgM, decreased after three subsequent vaccination injections Figure**a**. Furthermore, certain high-stability B-cell isotypes are found that offer the possibility of memory development and isotype flipping Figureb, c.

Additionally, during vaccination, the generation of memory cells (TCs) and CTL/HTL cells increased, suggesting immunogenicity in the presence of T-cell epitopes in the vaccine framework Figured-f. These findings generally indicate that the immune system experiences a constriction phase following the initial activation and multiplication of immune cells, which results in a drop in some immunoglobulins, such as IgM and IgG, between intervals following vaccination. This decrease in response is a typical aspect of the immune response and does not always signify vaccine failure. Other immune system elements, like memory T and B-cells, are crucial for preserving immunity in these circumstances.

Depending on the length of the infection’s incubation period, the quality of the memory response, and the quantities of memory B-cell antibodies, they enable quicker and more effective reactions when reexposed to future viruses. Consequently, the immune system is not helpless even if some antibody levels drop following the initial reaction (107). Additionally, dendritic cells (DCs) and macrophage activity increased with each exposure Figureg, h. A positive immune response is shown by high levels of IFN-γ, IL-23, IL-10, and IL-12, which dramatically rose following exposure Figurei. Lastly, immune simulations for three and twelve doses verify that the vaccine elicits a strong immune response and that the degree of protection rises even after repeated exposures in global settings.Additionally, dendritic cells (DCs) and macrophage activity increased with each exposure Figureg, h. A positive immune response is shown by high levels of IFN-γ, IL-23, IL-10, and IL-12, which dramatically rose following exposure Figurei. Lastly, immune simulations for three and twelve doses verify that the vaccine elicits a strong immune response and that the degree of protection rises even after repeated exposures in global settings. Antibodies, memory cells, and other immune responses are only a few of the immunological components that must be thoroughly evaluated in order to interpret vaccine efficacy and protection. Furthermore, the vaccine and the unique features of the immune system may affect the particular dynamics of the immune response (108). Supplementary Figure S18 offers more immunological simulation results

**Figure 11.**
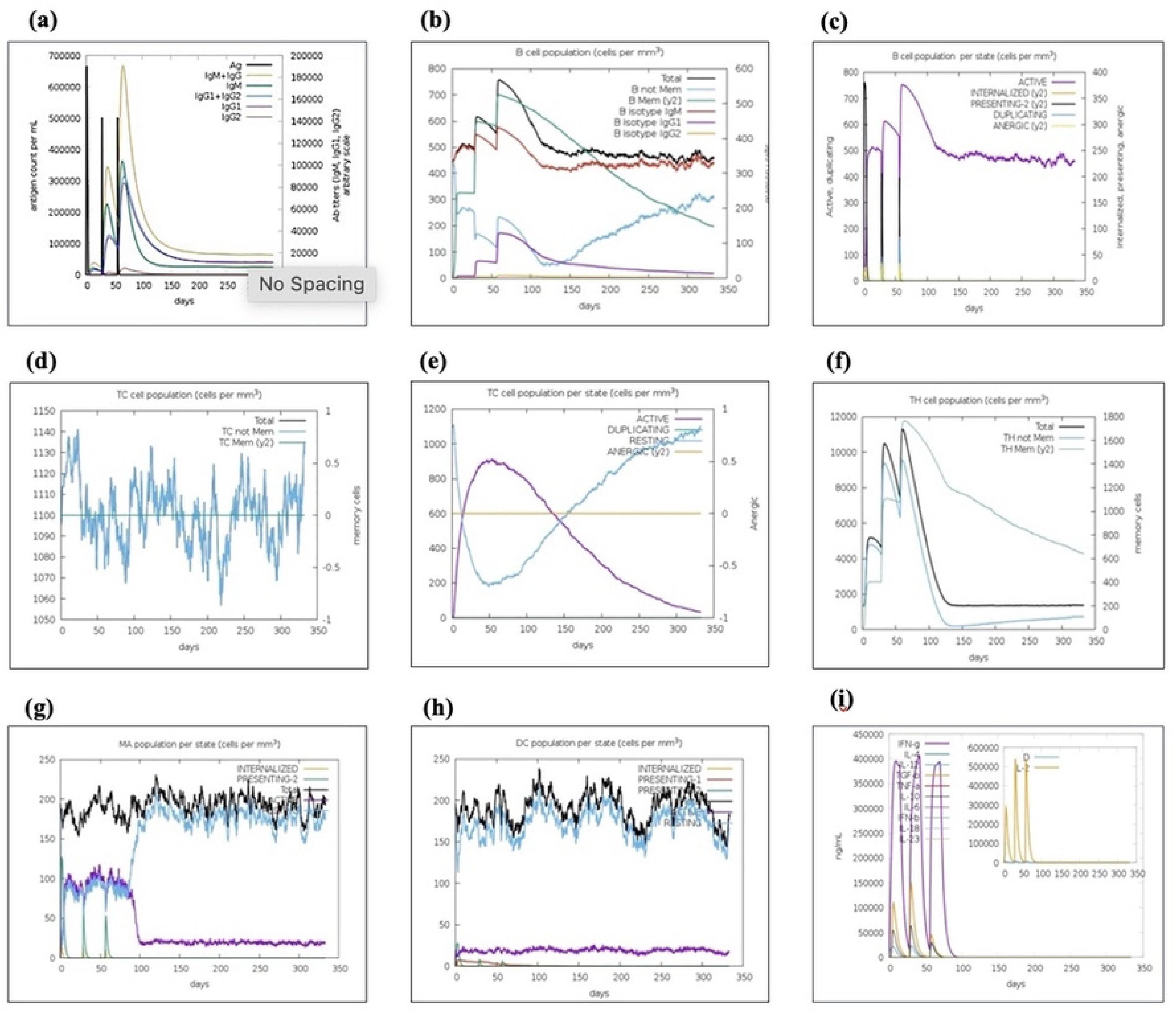
Shows an in-silico simulation of the immune response elicited by the vaccination as an antigen following three administrations. (a) Immunoglobulin subtypes and antigens are shown as distinct colored peaks. Following vaccination, the development of primary, secondary, and tertiary immune responses is reflected in the production of immunoglobulin (IgG). (b, c) The population and total count of B lymphocytes by entity state (active, presenting, internalized, replicating, or anergic). (d) The population of cytotoxic T cells. (e) The number of cytotoxic T cells in each state. (f) The population of helper T cells. (g) Population of macrophages by state. (h) Population of dendritic cells by state. (i) There are three stages in the concentration of interleukins and cytokines.

## Discussion

Helicobacter pylori remains a globally prevalent pathogen associated with chronic gastritis, peptic ulcer disease, and Helicobacter pylori is still a common infection in the world that is linked to stomach cancer, peptic ulcer disease, and chronic gastritis. Its persistence in the gastric mucosa, which is frequently formed in early childhood, presents a significant clinical problem. The effectiveness of traditional eradication therapy has been further hampered by the increasing appearance of antibiotic-resistant bacteria, highlighting the critical need for alternate prophylactic measures like immunization. In order to create a logical multi-epitope vaccine candidate that targets important H. pylori virulence-associated proteins, the current study used an integrated immunoinformatics and structural vaccinology method (109). The focused selection of adhesion-related and virulence-associated proteins, such as BabA, SabA, SabB, and VacA, which are essential for bacterial colonization, host contact, and persistence, is a distinguishing characteristic of this study.

Notably, this selection was guided by a pan-proteome-based approach, enabling the inclusion of conserved antigenic determinants across multiple H. pylori strains. By capturing proteomic diversity and prioritizing conserved epitopes, this strategy may enhance the breadth of immune coverage and increase the likelihood of cross-strain protection. While several previous immunoinformatics-based studies have designed multi-epitope vaccines using a broad range of antigenic proteins—such as CagA, urease subunits, OipA, and FliD—often aiming to maximize antigen diversity (110), the present study adopts a more focused and functionally relevant strategy by prioritizing proteins directly involved in host attachment and colonization. This targeted yet pan-proteome-informed design may improve the biological relevance of the elicited immune response by specifically interfering with key mechanisms underlying H. pylori persistence and pathogenicity (111).

In keeping with previous findings on multi-epitope vaccine design against H. pylori, the produced vaccine in this work demonstrated good antigenicity and structural features. For example, prior studies have demonstrated that multi-epitope constructions have intriguing docking interactions with immune receptors, stable structural conformations, and high antigenicity ratings (112). In a similar vein, the present construct demonstrated antigenicity scores of 0.49 (VaxiJen) and 0.84 (AntigenPro), indicating a strong potential for immune system recognition. However, the current study used a multi-parameter epitope selection pipeline that incorporates allergenicity, toxicity, conservation, population coverage, and non-homology to human proteins, increasing the likelihood of finding safe and widely applicable epitopes, in contrast to some previous designs that mainly rely on antigenicity thresholds. With an overall coverage of almost 92%, population coverage study provides additional evidence for the proposed vaccine’s potential worldwide applicability. This result is on par with or higher than that found in a number of earlier investigations, which generally fall between 80% and 90% depending on the methods used for epitope selection (109).

However, the comparatively lesser coverage shown in other areas, including Central America, draws attention to a significant drawback and implies that region-specific optimization would be necessary for the highest level of vaccine efficacy. Another crucial component of the vaccine design is the use of the cholera toxin B subunit (CTB) as a mucosal adjuvant. Prior research has shown that CTB promotes IgA-mediated immunity and facilitates antigen absorption through GM1 receptor binding to improve mucosal immune responses (112). According to these results, adding CTB to the current design might improve mucosal targeting, which is especially important considering the stomach niche of H. pylori. However, the new approach integrates CTB with a carefully chosen selection of adhesion-related epitopes, potentially leading to a more focused and functionally meaningful immune response, in contrast to previous research that mix CTB with a broad range of antigenic proteins.

Strong binding affinities between the vaccine construct and important immunological receptors, such as MHC class I (−15.9 kcal/mol), MHC class II (−27.1 kcal/mol), and TLR4 (−8.9 kcal/mol), were found by protein–protein interaction studies in this study. These results are in line with earlier studies showing stable docking connections between immune receptors and multi-epitope vaccines (110). Nonetheless, this study’s rather high interaction with MHC class II molecules points to a possible increase in CD4+ T helper cell activation, which is crucial for coordinating adaptive immune responses. Moreover, contact with TLR4 suggests that innate immune signaling pathways may be activated, resulting in the development of antigen-presenting cells and the release of pro-inflammatory cytokines.

The structural stability and adaptability of the vaccine construct under physiologically simulated settings were further validated by molecular dynamics simulations. The construct retains its conformational integrity throughout time, according to the comparatively stable RMSD profile and constant radius of gyration values (113). Previous research on multi-epitope vaccines has documented similar stability patterns, demonstrating the dependability of computational modeling techniques. According to RMSF analysis, the construct’s reported flexibility in some areas may promote epitope exposure and enhance immune recognition.

In silico immune simulations were used to further assess the vaccine’s immunogenic potential, and the results showed a coordinated and dynamic immune response. Class switching and memory development in classical immunological kinetics are consistent with the initial rise in IgM levels followed by increasing IgG subclasses in subsequent encounters. These results are consistent with earlier immunoinformatics research showing increased humoral responses after administering a multi-epitope vaccination (109). Notably, the current investigation also showed elevated numbers of helper T lymphocytes (HTLs) and cytotoxic T lymphocytes (CTLs), suggesting the activation of cellular immune responses.

Significantly, after immunization, cytokine profiling showed increased levels of IFN-γ, IL-12, IL-23, and IL-10. A robust Th1-biased immune response, which is crucial for managing intracellular infections like H. pylori, is suggested by the elevated IFN-γ levels. The precise correlation of IFN-γ with Th1-mediated protection in the context of H. pylori adds biological significance to the results of this work, even though other research has documented cytokine production in immunological simulations (112). A balanced immune response may be indicated by the concurrent rise in regulatory cytokines like IL-10, which may also aid in controlling excessive inflammation.

Despite these encouraging results, there are a few things to be aware of. First, the work relies solely on computer predictions, which might not adequately represent the intricacy of biological systems in vivo. Second, underlying algorithms and available datasets can bring bias or uncertainty into the predicted accuracy of immunoinformatics techniques. Third, experimental validation of the vaccine construct’s actual expression, folding, and stability in a biological system is still pending. The effectiveness of vaccines may also be impacted by environmental variables and host-specific immunological variability. Thus, additional in vitro and in vivo research is necessary to verify the immunogenicity, safety, and protective effectiveness of the suggested vaccination.

In summary, the current study offers a logical multi-epitope vaccination candidate against H. pylori that incorporates mucosal adjuvant inclusion, targeted antigen selection, and thorough computational evaluation. The results imply that the construct may maintain its structural and physicochemical characteristics while eliciting both humoral and cellular immune responses. The targeted selection of adhesion-related virulence factors in conjunction with CTB-mediated mucosal targeting is a potentially beneficial approach as compared to earlier research. However, to confirm these results and evaluate their potential for translation, experimental validation is needed.

## Conclusions

This work culminates in a logical multi-epitope vaccine candidate against Helicobacter pylori using an integrated immunoinformatics and pan-proteome-informed structural vaccinology strategy. By focusing on significant adhesion-related virulence proteins (BabA, SabA, SabB, and VacA), the construct was designed to target critical pathways of bacterial colonization and persistence, potentially enhancing functional immune interference against infection. A broader spectrum of H. pylori populations may be covered by the vaccine if conserved epitopes from a pan-proteome analysis are added.

Comprehensive in silico evaluations have shown that the proposed vaccine possesses optimal antigenicity, structural stability, and physicochemical properties as well as substantial binding affinity toward significant immunological receptors like MHC class I, MHC class II, and TLR4.Immunoglobulin class switching, memory cell development, and cytokine profiles associated with a Th1-biased immune response are characteristics of both humoral and cellular immune responses that may be induced, according to immunological simulation studies. This is especially pertinent for H. pylori clearance.

The current study takes a more functionally tailored and pan-proteome-informed approach than previously published multi-epitope vaccine designs, which frequently prioritize wide antigen inclusion. This approach may improve biological relevance and immunological specificity. Furthermore, the potential for increased stomach mucosal immunity—which is essential for fighting H. pylori infection—is further supported by the use of CTB as a mucosal adjuvant. The predicted immunological and structural characteristics, however, need experimental confirmation through in vitro and in vivo studies to confirm expression, safety, and protective efficacy because this work is fully computational. All things considered, this multi-epitope vaccine architecture is a logical and attractive option for future development against H. pylori infection, warranting further preclinical investigation.

## List of abbreviations

C-IMMSIM: Computational Immune Simulator
CAI: Codon adaptation index
CPORT: Consensus Prediction of Interface Residues in Transient complexes
CTB: Cholera toxin B subunit
GRAVY: Grand Average of Hydropathicity
Jcat: Java Codon Adaptation Tool
MALT: Mucosa-associated lymphoid tissue
MD: Molecular dynamics
PADRE: Pan DR epitope (universal helper T-cell epitope)
PDB: Protein Data Bank
PRODIGY: Protein binding affinity prediction server
Rg: Radius of gyration
RMSD: Root mean square deviation
RMSF: Root mean square fluctuation

## Declarations

### Ethics approval and consent to participate

Not applicable.

### Consent for publication

Not applicable.

### Availability of data and materials

The datasets generated and/or analyzed during the current study are available from the corresponding author upon reasonable request.

### Authors’ contributions

R.V. and N.H. contributed equally as co-first authors. R.V. and N.H. conceived and designed the study (Conceptualization). R.V. developed the immunoinformatics pipeline, performed molecular docking and molecular dynamics (MD) simulations, prepared the original draft (Methodology, Software, Investigation, Writing – original draft), and collected the H. pylori pan-proteome data, conducted the population coverage analysis, and prepared all figures including Supplementary Figure S2 (Data curation, Formal analysis, Visualization). R.A, N.A and A.M. participated in data interpretation, validated the computational results, and critically revised the manuscript (Validation, Writing – review & editing). All authors read and approved the final manuscript.

## Acknowledgements

The authors acknowledge the use of ChatGPT (OpenAI) for language editing and proofreading assistance during the preparation of this manuscript.

